# Adaptive stimulus selection for multi-alternative psychometric functions with lapses

**DOI:** 10.1101/260976

**Authors:** Ji Hyun Bak, Jonathan W. Pillow

## Abstract

Psychometric functions (PFs) quantify how external stimuli affect behavior and play an important role in building models of sensory and cognitive processes. Adaptive stimulus selection methods seek to select stimuli that are maximally informative about the PF given data observed so far in an experiment and thereby reduce the number of trials required to estimate the PF. Here we develop new adaptive stimulus selection methods for flexible PF models in tasks with two or more alternatives. We model the PF with a multinomial logistic regression mixture model that incorporates realistic aspects of psychophysical behavior, including lapses and multiple alternatives for the response. We propose an information-theoretic criterion for stimulus selection and develop computationally efficient methods for inference and stimulus selection based on semi-adaptive Markov Chain Monte Carlo (MCMC) sampling. We apply these methods to data from macaque monkeys performing a multi-alternative motion discrimination task, and show in simulated experiments that our method can achieve a substantial speed-up over random designs. These advances will reduce the data needed to build accurate models of multi-alternative PFs and can be extended to high-dimensional PFs that would be infeasible to characterize with standard methods.

## Introduction

Understanding the factors governing psychophysical behavior is a central problem in neuroscience and psychology. Although accurate quantification of the behavior is an important goal in itself, psychophysics provides an important tool for interrogating the mechanisms governing sensory and cognitive processing in the brain. As new technologies allow direct manipulations of neural activity in the brain, there is a growing need for methods that can characterize rapid changes in psychophysical behavior.

In a typical psychophysical experiment, an observer is trained to report judgements about a sensory stimulus by selecting a response from among two or more alternatives. The observer is assumed to have an internal probabilistic rule governing these decisions; this probabilistic map from stimulus to response is called the observer’s psychometric function. Because the psychometric function is not directly observable, it must be inferred from multiple observations of stimulus-response pairs. However, such experiments are costly due to the large numbers of trials typically required to obtain good estimates of psychometric functions. Therefore, a problem of major practical importance is to develop efficient experimental designs that can minimize the amount of data required to accurately infer an observer’s psychometric function.

### Bayesian adaptive stimulus selection

A powerful approach for improving the efficiency of psychophysical experiments is to design the data collection process so that the stimulus is adaptively selected on each trial by maximizing a suitably defined objective function (MacKay, 1992). Such methods are known by a variety of names, including “active learning”, “adaptive or sequential optimal experimental design”, and “closed-loop experiments.”

Bayesian approaches to adaptive stimulus selection define optimality of a stimulus in terms of its expected ability to improve the posterior distribution over the psychometric function, e.g., by reducing its variance or entropy. The three key ingredients of a Bayesian adaptive stimulus selection method are (Chaloner & Verdinelli, 1995; Pillow & Park, 2016):

- **model** - parametrizes the psychometric function of interest;
- **prior** - captures initial beliefs about model parameters;
- **utility function** - quantifies the usefulness of a hypothetical stimulus-response pair for improving the posterior.

Sequential algorithms for adaptive Bayesian experiments rely on repeated application of three basic steps: (i) data collection (stimulus presentation and response measurement); (ii) inference (posterior updating using data from the most recent trial); and (iii) selection of an optimal stimulus for the next trial by maximizing expected utility (see Fig. 1A). The inference step involves updating the posterior distribution over the model parameters according to Bayes rule with data from the most recent trial. Stimulus selection involves calculating the expected utlity (i.e., the expected improvement in the posterior) for a set of candidate stimuli, averaging over the responses that might be elicited for each stimulus, and selecting the stimulus for which the expected utility is highest. Example utility functions include the negative trace of the posterior covariance (corresponding to the sum of the posterior variances for each parameter) and the mutual information or information gain (which corresponds to minimizing the entropy of the posterior).

Methods for Bayesian adaptive stimulus selection have been developed over several decades in a variety of different disciplines. If we focus on the specific application of estimating psychometric functions, the field goes back to the QUEST (A. B. Watson & Pelli, 1983) and ZEST (King-Smith, Grigsby, Vingrys, Benes, & Supowit, 1994) algorithms, which were focused on the estimation of discrimination thresholds, and to the simple case of 1-dimension stimulus and binary responses (Treutwein, 1995). The Ψ method (Kontsevich & Tyler, 1999) used Bayesian inference for estimating both threshold and slope of a psychometric function, which were extended to two-dimensional stimuli by Kujala and Lukka (2006). Further development of the method allowed for adaptive estimation of more complex psychometric functions, where the parameters were no longer limited to a threshold and a slope (Barthelmé & Mamassian, 2008; Kujala & Lukka, 2006; Lesmes, Lu, Baek, & Albright, 2010; Prins, 2013); and possibly related to each other (Vul, Bergsma, & MacLeod, 2010). Models with multidimensional stimuli were also considered (DiMattina, 2015; Kujala & Lukka, 2006; A. B. Watson, 2017).

Different models have been used to describe the psychometric function. Standard choices include the logistic regression model (Chaloner & Larntz, 1989; DiMattina, 2015; Zocchi & Atkinson, 1999), the Weibull distribution function (A. B. Watson & Pelli, 1983), and the cumulative function of Gaussian distribution (Kontsevich & Tyler, 1999). More recent works also considered Gaussian Process models (Gardner, Song, Weinberger, Barbour, & Cunningham, 2015). Most of the previous works, however, were limited to the case of binary responses.

In parallel, the development of Bayesian methods for inferring psychometric functions (Kuss, Jäkel, & Wichmann, 2005; Prins, 2012; Wichmann & Hill, 2001a, 2001b) have enlarged the space of statistical models that could be employed to describepsychophys-ical phenomena based on (often limited) data. A variety of recent advances also arose in sensory neuroscience or neurophysiology, driven by the development of efficient inference techniques for neural encoding models (Lewi, Butera, & Paninski, 2009; M. Park, Horwitz, & Pillow, 2011) or model comparison and discrimination methods (Cavagnaro, Myung, Pitt, & Kujala, 2010; DiMattina & Zhang, 2011; Kim, Pitt, Lu, Steyvers, & Myung, 2014). These advances can in many cases be equally well applied to psychophysical experiments.

### Our contributions

In this paper, we develop methods for adaptive stimulus selection in psychophysical experiments that are applicable to realistic models of human and animal psychophysical behavior. First, we describe a psychophysical model that incorporates multiple response alternatives and “lapses”, in which the observer makes a response that does not depend on the stimulus. This model can also incorporate “omission” trials, where the observer does not make a valid response (e.g., breaking fixation before the go cue), by considering them as an additional response category. Second, we describe efficient methods for updating the posterior over the model parameters after every trial. Third, we introduce two algorithms for adaptive stimulus selection, one based on a Gaussian approximation to the posterior and a second based on Markov Chain Monte Carlo (MCMC) sampling. We apply these algorithms to simulated data and to real data analyzed with simulated closed-loop experiments, and show that they can substantially reduce in the number of trials required to estimate multi-alternative psychophysical functions.

**Figure 1:**
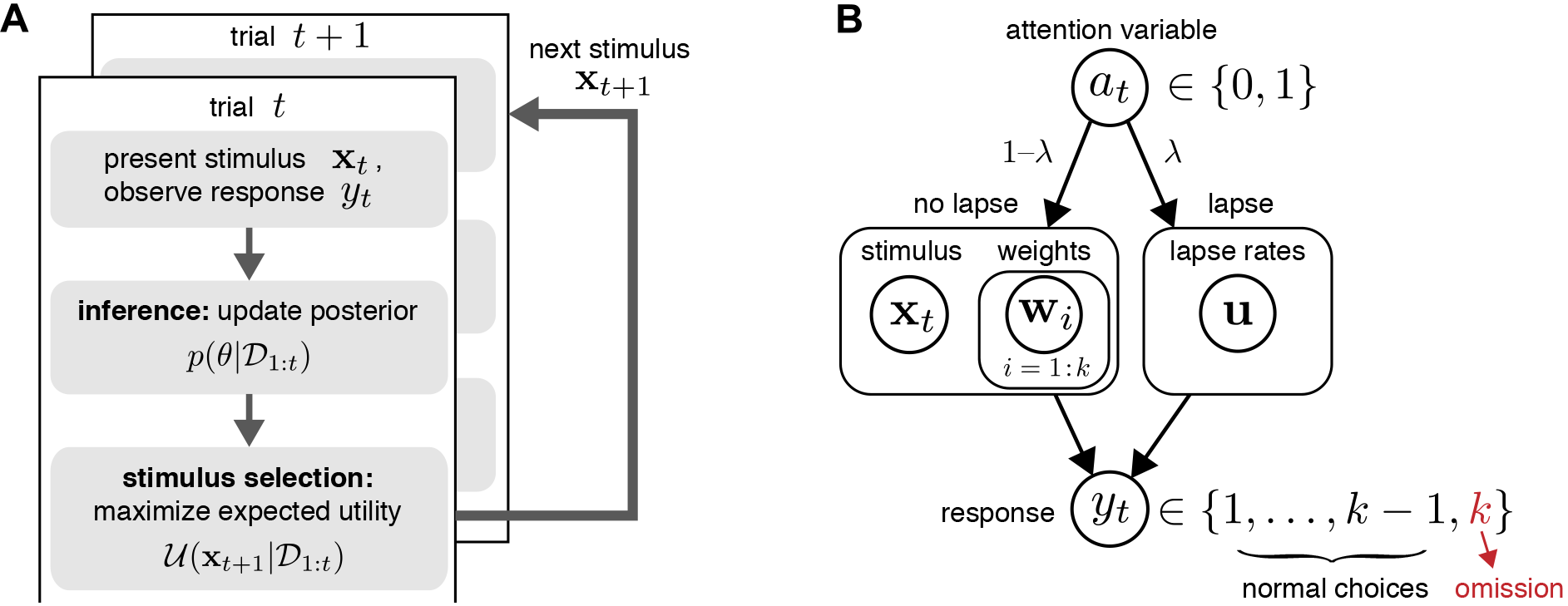
**(A)** Schematic of Bayesian adaptive stimulus selection. On each trial: (i) a stimulus is presented and response is observed; (ii) the posterior over the parameters *θ* is updated using all data collected so far in the experiment 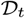; and (iii) the stimulus that maximizes the expected utility (in our case, information gain) is selected for the next trial. **(B)** A graphical model illustrating a hierarchical psychophysical observer model that incorporates lapses as well as the possibility of omissions. On each trial, a latent attention or lapse variable *a*_*t*_ is drawn from a Bernoulli distribution with parameter λ, to determine whether the observer attends to the stimulus **x**_*t*_ on that trial or lapses. With probability 1 − λ, and the observer attends to the stimulus (*a*_*t*_ = 0), and the response *y*_*t*_ is drawn from a multinomial logistic regression model, where the probability of choosing option *i* is proportional to 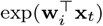. With probability λ, the observer lapses (*a*_*t*_ = 1) and selects a choice from a (stimulus-independent) response distribution governed by parameter vector **u**. So-called “omission” trials, in which the observer does not select one of the valid response options, are modeled with an additional response category *y*_*t*_ = *k*.

## Psychophysical observer model

Here we describe a flexible model of psychometric functions (PFs) based on the multinomial logistic (MNL) response model (Glonek & McCullagh, 1995). We show how omission trials can be naturally incorporated into a model as one of the multiple responses alternatives. We then develop a hierarchical extension of the model that incorporates lapses (see Fig. 1B).

### Multinomial logistic response model

We consider the setting where the observer is presented with a stimulus x ∈ ℝ^*d*^ and selects a response *y* ∈ {1,… *k*} from one of *k* discrete choices on each trial. We will assume the stimulus is represented internally by some (possibly non-linear) feature vector *ϕ*(x), which we will write simply as *ϕ* for notational simplicity.

In the multinomial logistic model, the probability *p*_*i*_ of each possible outcome *i* ∈ {1, ···, *k*} is determined by the dot product between the feature *ϕ* and a vector of weights w_*i*_ according to:

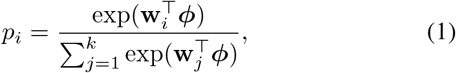

where the denominator ensures that these probabilities sum to 1, 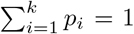. The function from stimulus to a probability vector over choices, x ↦ (*p*_1_,… *p*_*k*_), is the psychometric function, and the set of weights 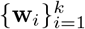 are its parameters. Note that the model is over-parameterized when written this way, since the requirement that probabilities sum to 1 removes one degree of freedom from the probability vector. Thus, we can without loss of generality fix one of the weight vectors to zero, for example w_*k*_ = **0**, so that the denominator in (eq. 1) becomes 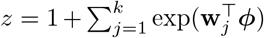 and *P*_*k*_ = 1/*z*.

We consider the feature vector *ϕ* to be a known function of the stimulus x, even when the dependence is not written explicitly. For example, we can consider a simple form of feature embedding, *ϕ*(x) = [1, x^⊤^]^⊤^, corresponding to a linear function of the stimulus plus an offset. In this case, the weights for the *i*’th choice would correpond to 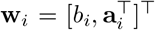, where *b*_*i*_ is the offset or bias for the *i*’th choice, and ***a***_*i*_ are the linear weights governing sensitivity to x. The resulting choice probability has the familiar form, 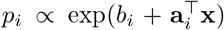. Nonlinear stimulus dependencies can be incorporated by including nonlinear functions of x in the feature vector *ϕ*(x) (Knoblauch & Maloney, 2008; Murray, 2011; Neri & Heeger, 2002). Dependencies on the trial history, such as the previous stimulus or reward, may also be included as additional features in *ϕ* (see for example Bak, Choi, Akrami, Witten, and Pillow (2016)).

It is useful to always work with a normalized stimulus space, in which the mean of each stimulus component *x*_*α*_ over the stimulus space is 〈*x*_*α*_〉 = 0, and the standard deviation std(*x*_*α*_) = 1. This normalization ensures that the values of the weight parameters are defined in more interpretable ways. The zero-mean condition ensures that the bias *b* is the expectation value of log probability over all possible stimuli. The unit-variance condition means that the effect of moving a certain distance along one dimension of the weight space is comparable to the moving the same distance in another dimension, again averaged over all possible stimuli. In other words, we are justified to use the same unit along all dimensions of the weight space.

### Including omission trials

Even in binary tasks with only two possible choices per trial, there is often an implicit third choice, which is to make no response, make an illegal response, or to interrupt the trial at some point before the allowed response period. For example, animals are often required to maintain an eye position or a nose poke, or wait for a “go” cue before reporting a choice. Trials on which the animal fails to obey these instructions are commonly referred to as “omissions” or “violations”, and are typically discarded from analysis. However, failure to take these trials into account may bias the estimates of the PF if they are more common for some stimuli than others (see Fig. 2B).

The multinomial response model provides a natural framework for incorporating omission trials because it accommodates an arbitrary number of response categories. Thus we can model omissions explicitly as one of the possible choices the observer can choose from, or as the (*k* + 1)’st response category in addition to the *k* valid responses. One could even consider different kinds of omissions separately, e.g., allowing choice *k*+1 to reflect fixation period violations and choice *k* + 2 to reflect failure to report a choice during the response window. Henceforth, we will let *k* reflect the total number of choices, including omission, as illustrated in Fig. 1B.

This formulation can also be useful for the rated Yes/No task in human psychophysics, where a “Not Sure” response is explicitly presented (C. S. Watson, Kellogg, Kawanishi, & Lucas, 1973). Although such model was considered for adaptive stimulus selection (Lesmes et al., 2015), the third alternative was not handled as a fully independent choice, as the goal was only to estimate the two detection thresholds separately: one for a strict Yes, another for a collapsed response of either Yes or Not Sure. Our model treats each of the multiple alternatives equivalently.

**Figure 2:**
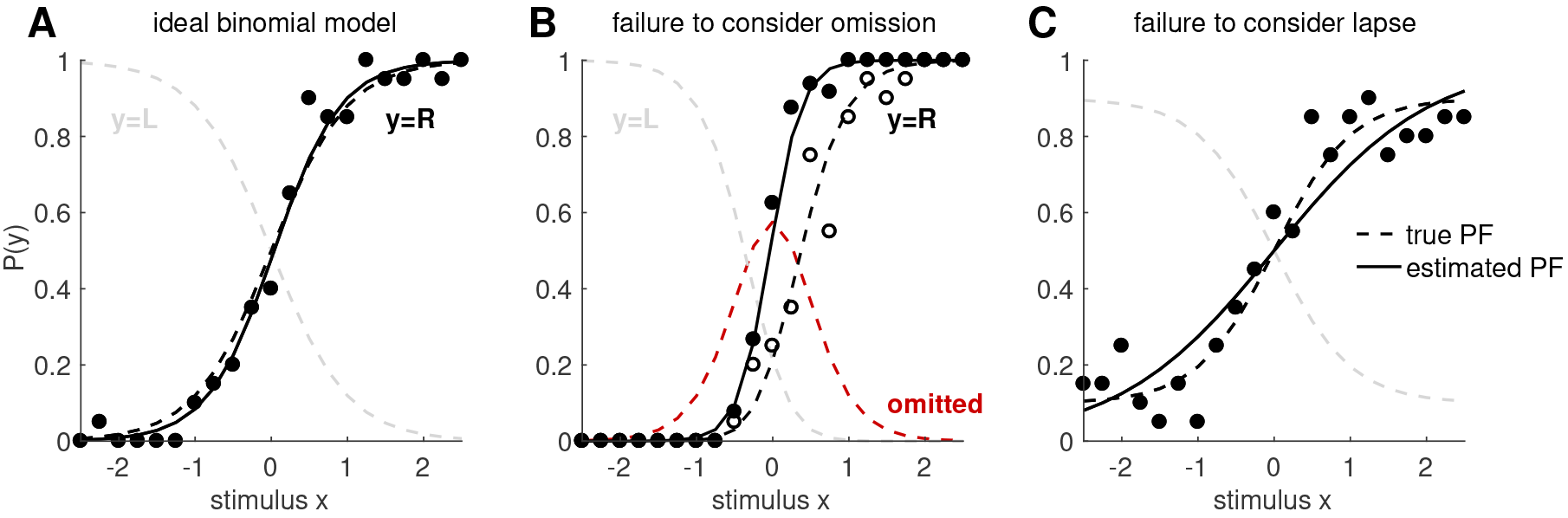
Effects of omission and lapse. Here we illustrate the undesirable effects of failing to take into account of omission and lapse. **(A)** If the PF follows an ideal binomial logistic model, it can be estimated very well from data. The black dashed line shows the true PF for one of the two responses (say *y* = *R*), and the gray dashed line shows the true PF for the other response (say *y* = *L*), such that the two dashed curves always add up to 1. The black dots indicate the mean probability to observe this response *y* = *R* at each stimulus point *x*. We drew 20 observations per stimulus point, at each of the 21 stimulus points along the 1-dimensional axis. The resulting estimate for *P*(*y* = 1) is shown in the solid black line. The inference method is not important for the current purpose, but we used the MAP estimate, discussed in a later section. **(B)** Now suppose that some trials fell into the implicit third choice which is the omission (red dashed line shows omission probability). The observed probability of *y* = *R* at each stimulus point (open black circles) follows the true PF (black dashed line). But if the omitted trials are systematically excluded from analysis, as in common practice, the estimated PF (solid black line) reflects a biased set of observations (filled black circles), and fail to recover the true PF. **(C)** When there is a finite lapse rate (we used a total lapse of λ = 0:2, uniformly distributed to the two outcomes), the true PF (dashed black line) asymptotes to a finite offset from 0 or 1. If the resulting observations (black dots) are fitted to a plain binomial model without lapse, the slope of the estimated PF (solid black line) is systematically biased.

### Modeling lapse with a mixture model

Another important feature of real psychophysical observers is the tendency to occasionally make errors that are independent of the stimulus. Such choices, commonly known as “lapses” or “button press errors”, may reflect lapses in concentration or memory of the response mapping (Kuss et al., 2005; Wichmann & Hill, 2001a). Lapses are most easily identified by errors on “easy” trials, that is, trials that should be performed perfectly if the observer were paying attention.

Although lapse rates can be negligible in highly trained observers (Carandini & Churchland, 2013), they can be substantially greater than zero in settings involving non-primates or complicated psychophysical tasks. Lapses affect the psychometric function by causing it to saturate above 0 and below 1, so that “perfect” performance is never achieved even for the easiest trials. Failure to incorporate lapses into the PF model may therefore bias estimates of sensitivity, as quantified by PF slope or threshold (illustrated in Fig.2C; also see Wichmann and Hill (2001a, 2001b) or Prins (2012)).

To model lapses, we use a mixture model that treats the observer’s choice on each trial as coming from one of two probability distributions: a stimulus-dependent distribution (governed by the multinomial logistic model) and stimulus-independent distribution (reflecting a fixed probability of choosing any option when lapsing). Simpler versions of such mixture model have been proposed previously (Kuss et al., 2005).

Fig. 1B shows a schematic of the resulting model. On each trial, a Bernoulli random variable *a* ~ Ber(λ) governs whether the observer lapses: with probability λ and the observer lapses (i.e., ignores the stimulus), and with probability 1 − λ, and the observer attends to the stimulus. If the observer lapses (*a* = 1), the response is drawn according to fixed probability distribution (*c*_1_,…,*c*_*k*_) governing the probability of selecting options 1 to *k*, where ∑*C*_*i*_ = 1. If the observer does not lapse (*a* = 0), the observer selects a re-sponse according to the multinomial logistic model. Under this model, the conditional probability of choosing option *i* given the stimulus can be written:

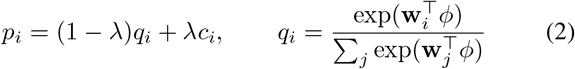

where *q*_*i*_ is the lapse-free probability probability under the classical MNL model (eq. 1).

It is convenient to re-parameterize this model so that *λc*_*i*_, the conditional probability of choosing the *i*-th option due to a lapse, is written

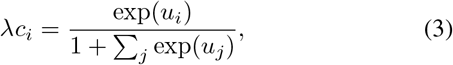

where each auxiliary lapse parameter *u*_*i*_ is proportional to the log probability of choosing option *i* due to lapse. The lapse-conditional probabilities of each choice, *c*_*i*_, and the total lapse probability, λ, are respectively

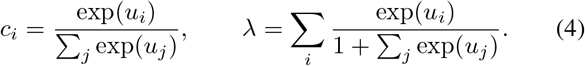

Because each *u*_*i*_ lives on the entire real line, fitting can be carried out with unconstrained optimization methods, although adding reasonable constraints may improve performance in some cases. The full parameter vector of the resulting model is *θ* = [**w**^⊤^, **u**^⊤^]^⊤^, which includes *k* additional lapse parameters **u** = {*u*_1_, · · ·, *u*_*k*_}. Note that in some cases it might be desirable to assume lapse choices obey a uniform distribution, where the probability of each option is *c*_*i*_ = 1/*k*. For this simplified “uniform-lapse” model we need only a single lapse parameter *u*. Note that we have unified the parameterizations of the “lapse rate” (deviation of the upper asymptote of the PF from 1; in this case λ - λ*c*_*i*_) and the “guess rate” (deviation of the lower asymptote from 0; in this case λ*c*_*i*_), which are often modeled separately in previous works with two-alternatives responses (Schutt, Harmeling, Macke, & Wichmann, 2016; Wichmann & Hill, 2001a, 2001b). Here they are written in terms of a single family of parameters {*u*_*i*_}, and extended naturally to multi-alternative responses.

Our model provides a general and practical parametrization of psychometric functions with lapses. Although previous work has considered the problem of modeling lapses in psychophysical data, much of it assumed a uniform-lapse model, where all options are equally likely during lapses. Earlier approaches have often assumed either that the lapse probability was known a priori (Kontsevich & Tyler, 1999), or was fit by a grid search over a small set of candidate values (Wichmann & Hill, 2001a). Here we instead infer individual lapse probabilities for each response option, similar to recent approaches described in Kuss et al. (2005); Prins (2012, 2013); Schütt et al. (2016). Importantly, our method infers the full parameter 0 that includes both the weight and the lapse parameters, rather than treating the lapse separately. In particular, our parameterization (eq. 3) has the advantage that there is no need to constrain the support of the lapse parameters *u*_*i*_. These parameters’ relationship to lapse probabilities *c*_*i*_ takes the same (“softmax”) functional form as the multinomial logistic model, placing both sets of parameters on an equal footing.

Before closing this section, we would like to reflect briefly on the key differences between omissions and lapses. First, although omissions and lapses both reflect errors in decision making, omissions are defined as invalid responses and are thus easily identifiable from the data; lapses, on the other hand, are indistinguishable from normal responses, and are identifiable only from the fact that the psychometric function does not saturate at 0 or 1. Second, modeling omissions as a response category under the multinomial logistic model means that the probability of omission is stimulus-dependent (e.g., more likely to arise on trials with high difficulty, or generally when the evidence for other options is low). Even if the omissions are not stimulus-dependent, and governed entirely by a “bias” parameter, the probability of omission will still be higher when the evidence for other choices is low, or lower when the evidence for other choices is high. Omissions that arise in a purely stimulus-independent fashion, on the other hand, will be modeled as arising from the lapse parameter associated with the omission response category. Omissions can thus arise in two ways under the model: as categories selected under the multinomial model or as lapses arising independent of the stimulus and other covariates.

## Posterior inference

Bayesian methods for adaptive stimulus selection require the posterior distribution over model parameters given the data observed so far in an experiment. The posterior distribution results from the combination of two ingredients: a prior distribution *p*(***θ***), which captures prior uncertainty about the model parameters ***θ***, and a likelihood function *p*({*y*_*s*_}|{x_*s*_}, ***θ***), which captures information about the parameters from the data {(x_*s*_, *y*_*s*_)}, *s* = 1,…, *t*, consisting of stimulus-response pairs observed up to the current time bin *t*.

Unfortunately, the posterior distribution for our model has no analytic form. We therefore describe two methods for approximate posterior inference: one relying on a Gaussian approximation to the posterior, known as the Laplace approximation, and a second one based on MCMC sampling.

### Prior

The prior distribution specifies our beliefs about model parameters before we have collected any data, and serves to regularize estimates obtained from small amounts of data, e.g., by shrinking estimated weights toward zero. Typically we want the prior to be weak enough that the likelihood dominates the posterior for reasonable-sized datasets. However, the choice of prior is especially important in adaptive stimulus selection settings because it determines the effective volume of the search space (M. Park & Pillow, 2012; M. Park, Weller, Horwitz, & Pillow, 2014). For example, if the weights are known to exhibit smoothness, then a correlated or smoothness-inducing prior can improve the performance of adaptive stimulus selection because the effective size (or entropy) of the parameter space is much smaller than under an independent prior (M. Park & Pillow, 2012).

In this study, we use a generic independent, zero-mean Gaussian prior over the weight vectors

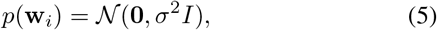

for all *i* ∊ (1,… *k*), with a fixed standard deviation *σ*. This choice of prior is appropriate when the regressors {x} are standardized, since any single weight can take values that allow for a range of psychometric function shapes along that axis, from flat (*w* = 0) to steeply decreasing (*w* = −2*σ*) or increasing (*w* = +2*σ*). We used *σ* = 3 in the simulated experiments in Results. For the lapse parameters {*u*_*i*_}, we used a uniform prior over the range [log(0.001), 0] with the natural log, so that each lapse probability λc_*i*_ is bounded between 0.001 and 1/2. We set the lower range constraint below 1/*N*, where *N* = 100 is the number of observed trials in our simulations, since we cannot reasonably infer lapse probabilities with precision finer than 1/*N*. The upper range constraint gives maximal lapse probabilities of 1/(*k* +1) if all *u*_*i*_ take on the maximal value of 0. Note that our prior is uniform with respect to the rescaled lapse parameters {*u*_*i*_}, rather than to the actual lapse rates; projected to the space of the lapse probabilities, given the bounds, the prior increases towards smaller lapse. For a comprehensive study of the effect of different priors on lapse, see for example Schütt et al. (2016).

### Psychometric function likelihood

The likelihood is the conditional probability of the data as a function of the model parameters. Although we have thus far considered the response variable *y* to be a scalar taking values in the set {1,…, *k*}, it is more convenient to use a “one-hot” or “1-of-*k*” representation, in which the response variable y for each trial is a length-*k* vector with one 1 and (*k* - 1) zeros; the position of the 1 in this vector indicates the category selected. For example, in a task with four possible options per trial, a response vector y =[0010] indicates a trial on which the observer selected the third option.

With this parametrization, the log-likelihood function for a single trial can be written

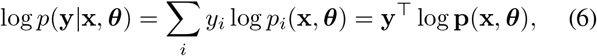

where *p*_*i*_(x, ***θ***) denotes the probability *p*(*y*_*i*_ = 1|x, ***θ***) under the model (eq. 1), and **p**(x, ***θ***) ≡ [*p*_1_(x, ***θ***),…, *p*_*k*_(x, ***θ***)]^⊤^ denotes the vector of probabilities for a single trial.

In the classical (lapse-free) multinomial logistic model, where ***θ*** = {*w*_*i*_}, the log likelihood is a concave function of ***θ***, which guarantees that numerical optimization of the log-likelihood will find a global optimum. With a finite lapse rate, however, the log likelihood is no longer concave. (See Appendix A).

### Posterior distribution

The log-posterior can be written as the sum of log-prior and log-likelihood summed over trials, plus a constant:

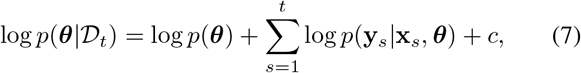

where 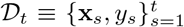 denotes the accumulated data up to trial *t* and *c* = − log (∫ *p*(***θ***) Π_*s*_ *p*(**y**_*s*_ |**x**_*s*_)*d****θ***) is a normalization constant that does not depend on the parameters *θ*. Because this constant has no tractable analytic form, we rely on two alternate methods for obtaining a normalized posterior distribution.

**Inference via Laplace approximation.** The Laplace approximation is a well-known Gaussian approximation to the posterior distribution, which can be derived from a second-order Tayler series approximation to the log-posterior around its mode (Bishop, 2006).

Computing the Laplace approximation involves a two-step procedure. The first step is to perform a numerical optimization of 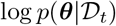 to find the posterior mode, or maximum a posteriori (MAP) estimate of *θ*. This vector, given by

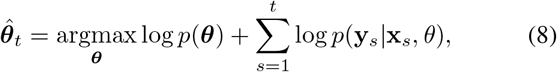

provides the mean of the Laplace approximation. Because we can explicitly provide the gradient and Hessian of the log likelihood (see Appendix A) and log-prior, this optimization can be carried efficiently via Newton-Raphson or trust region methods.

The second step is to compute the second derivative (the Hessian matrix) of the log-posterior at the mode, which provides the inverse covariance of the Gaussian. This gives us a local Gaussian approximation of the posterior, centered at the posterior mode:

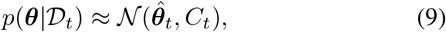

where covariance 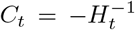 is the inverse Hessian of the log posterior, 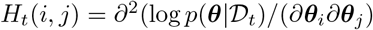, evaluated at 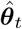.

Note that when the log-posterior is concave (i.e., when the model does *not* contain lapse), numerical optimization is guaranteed to find a global maximum of the posterior. Log-concavity also strengthens the rationale for using the Laplace approximation, since the true and approximate posterior are both log-concave densities centered on the true mode (Paninski et al., 2010; Pillow, Ah-madian, & Paninski, 2011). When the model incorporates lapses, these guarantees no longer apply globally.

### Inference via MCMC sampling

A second approach to inference is to generate samples from the posterior distribution over the parameters via Markov Chain Monte Carlo (MCMC) sampling. Sampling-based methods are typically more computationally intensive than the Laplace approximation, but may be warranted when the posterior is not provably log-concave (as is the case when lapse rates are non-zero) and therefore not well approximated by a single Gaussian.

The basic idea in MCMC sampling is to set up an easy-to-sample Markov Chain that has the posterior as its stationary distribution. Sampling from this chain produces a dependent sequence of posterior samples: 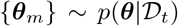, which can be used to evaluate posterior expectations via Monte Carlo integrals:

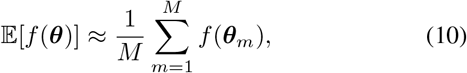

for any function *f*(***θ***). The mean of the posterior is obtained from setting *f*(***θ***) = ***θ***, although for adaptive stimulus selection we will be interested in the full shape of the posterior.

The Metropolis-Hastings (MH) algorithm is perhaps the simplest and most widely-used MCMC sampling method (Metropolis, Rosenbluth, Rosenbluth, Teller, & Teller, 1953). It generates samples via a proposal distribution centered on the current sample (see Appendix B). The choice of proposal distribution is critical to the efficiency of the MH algorithm, since this governs the rate of “mixing”, or the the number of Markov Chain samples required to obtain independent samples from the posterior distribution (Rosenthal, 2011). Faster mixing implies that fewer samples *M* are required to obtain an accurate approximation to the posterior.

**Figure 3:**
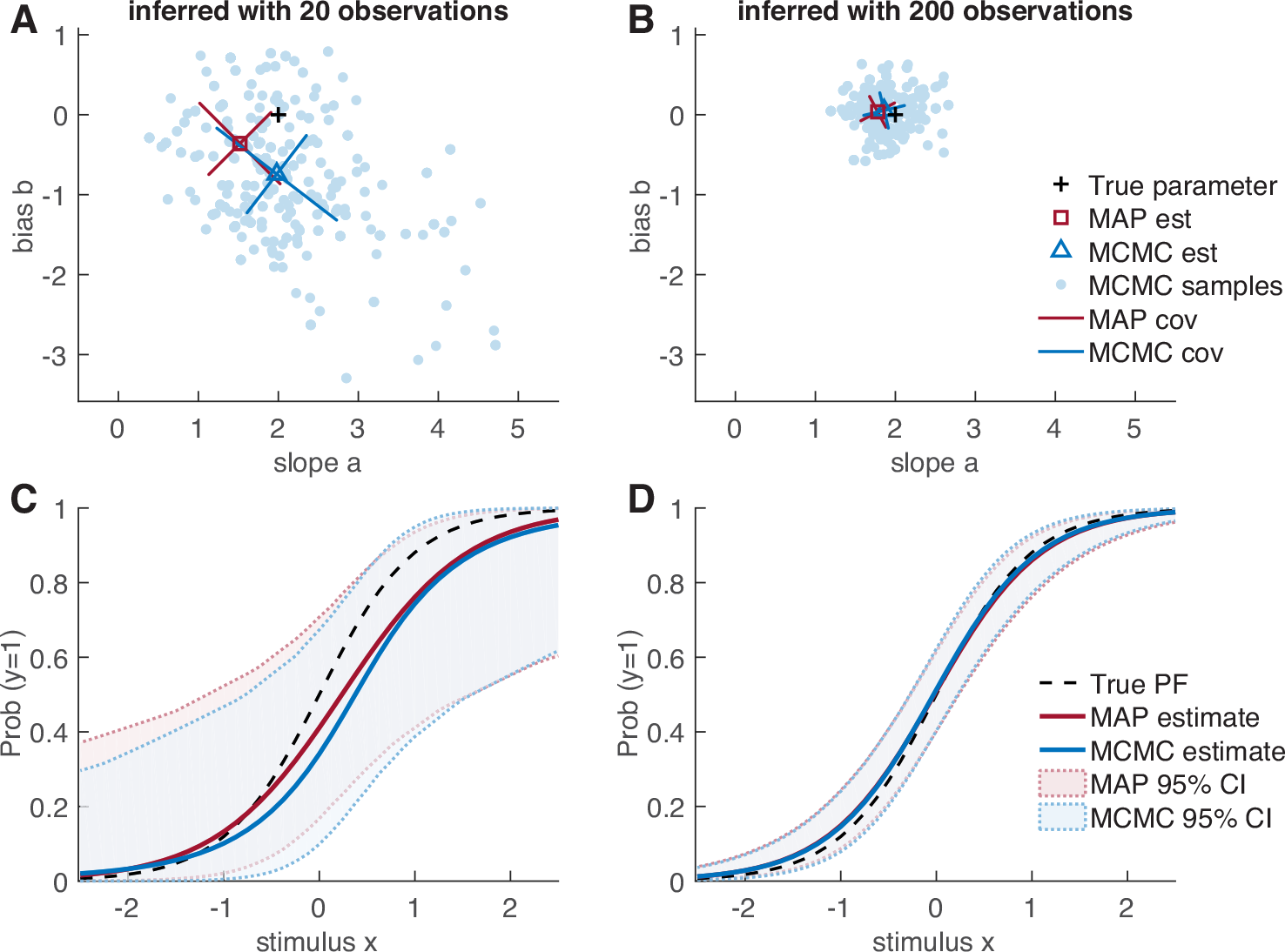
Inferring the psychometric function. Example of a psychometric problem, with a lapse-free binomial logistic model *f*(*v*) = *e*^*v*^=/(1+*e*_*v*_). Given a 1D stimulus, a response were drawn from a “true” model *P*(*y* = 1) = *f*(*b* + *ax*) with two parameters, slope *a* = 2 and bias *b* = 0. **(A-B)** Viewing on the parameter space, the posterior distributions become sharper (and closer to the true parameter values) as the dataset size *N* increases. Shown at a small **(A)** *N* = 20 and a large **(B)** *N* = 200. For the MAP estimate, the mode of the distribution is marked with a square, and the two standard deviations (“widths”) of its Gaussian approximation are shown with bars. For the MCMC sampling method, all *M* = 500 samples of the chain are shown in dots, the sample mean with a triangle, and the widths with the bars. The widths are the standard deviations along the principal directions of the sampled posterior (eigenvectors of the covariance matrix; not necessary aligned with the a − b axes). **(C-D)** The accuracy of the estimated PF improves with the number of observations *N*, using either of the two posterior inference methods (MAP-based and sampling-based). Shown at a small **(C)** *N* = 20 and a large **(D)** *N* = 200. The two methods are highly consistent in this simple case, especially when *N* is large enough.

Here we propose a semi-adaptive MH algorithm, developed specifically for the current context of sequential learning. Our approach is based on an established observation that the optimal width of the proposal distribution should be proportional to the typical length scale of the distribution being sampled (Gelman, Roberts, & Gilks, 1996; Roberts, Gelman, & Gilks, 1997). Our algorithm is motivated by the adaptive Metropolis algorithm (Haario, Saksman, & Tamminen, 2001), where the proposal distribution is updated at each proposal within a single chain; here we do not adapt the proposal within chains, but rather after each trial. Specifically, we set the covariance of a Gaussian proposal distribution to be proportional to the covariance of the samples from the previous trial, using the scaling factor of Haario et al. (2001). See Appendix B for details. The adaptive algorithm takes advantage of the fact that the posterior cannot change too much between trials, since it changes only by a single-trial likelihood term on each trial.

## Adaptive stimulus selection methods

As data are collected during the experiment, the posterior distribution becomes narrower due to the fact that each trial carries some additional information about the model parameters (see Fig. 3). This narrowing of the posterior is directly related to information gain. A stimulus that produces no expected narrowing of the posterior is, by definition, uninformative about the parameters. On the other hand, a stimulus that (on average) produces a large change in the current posterior is an informative stimulus. Selecting informative stimuli will reduce the number of stimuli required to obtain a narrow posterior, which is the essence of adaptive stimulus selection methods. In this section, we introduce a precise measure of information gain between a stimulus and the model parameters, and propose an algorithm for selecting stimuli to maximize it.

### Infomax criterion for stimulus selection

At each trial, we present a stimulus x and observe the outcome y. After *t* trials, the expected gain in information from a stimulus x is equal to the mutual information between y and the model parameters ***θ***, given the data 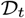 observed so far in the experiment. We denote this conditional mutual information:

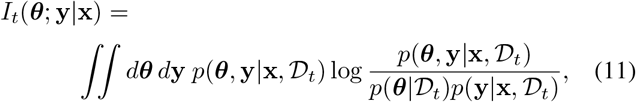

where 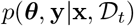 is the joint distribution of ***θ*** and y given a stimulus x and dataset 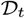, the term 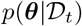 is the current posterior distribution over the parameters from previous trials, and 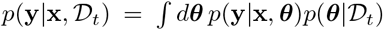 is known as the posterior-predictive distribution of y given x.

It is useful to note that the mutual information can equivalently be written in two other ways involving Shannon entropy. The first is given by:

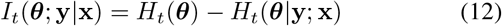

where the first term is the entropy of the posterior at time *t*,

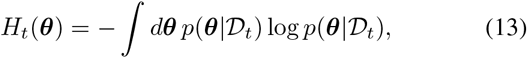

and the second is the conditional entropy of ***θ*** given y,

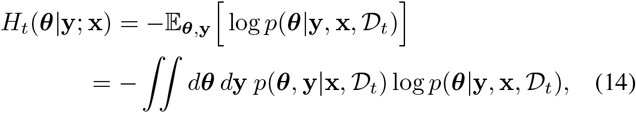

which is the entropy of the updated posterior *after* having observed x and y, averaged over draws of y from the posterior predictive distribution. Written this way, the mutual information can be seen as the expected reduction in posterior entropy from a new stimulus-response pair. Moreover, the first term, *H*_*t*_(*θ*), is independent of the stimulus and response on the current trial, so infomax stimulus selection is equivalent to picking the stimulus that minimizes the expected posterior entropy *H*_*t*_ (***θ***|y; x).

A second equivalent expression for the mutual information, which will prove useful for our sampling-based method, is:

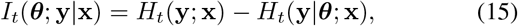

which is the difference between the marginal entropy of the response distribution conditioned on x,

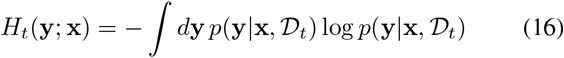

and the conditional entropy of the response y given ***θ***, conditioned on the stimulus:

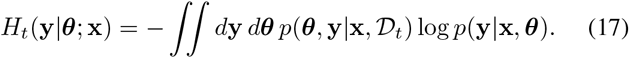

This formulation shows the mutual information to be equal to the difference between the entropy of the marginal distribution of y conditioned on x (with ***θ*** integrated out) and the average entropy of y given x and ***θ***, averaged over the posterior distribution of ***θ***. The dual expansion of the mutual information was also used by Kujala and Lukka (2006).

In a sequential setting where *t* is the latest trial and *t* + 1 is the upcoming one, the optimal stimulus is the information-maximizing (“infomax”) solution:

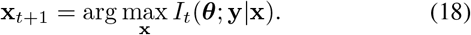

Fig. 4 shows an example of a simulated experiment where the stimulus was selected adaptively following the infomax criterion. Note that our algorithm takes a “greedy” approach of optimizing one trial at a time. For work on optimizing beyond the next trial, see for example Kim, Pitt, Lu, and Myung (2017).

**Figure 4:**
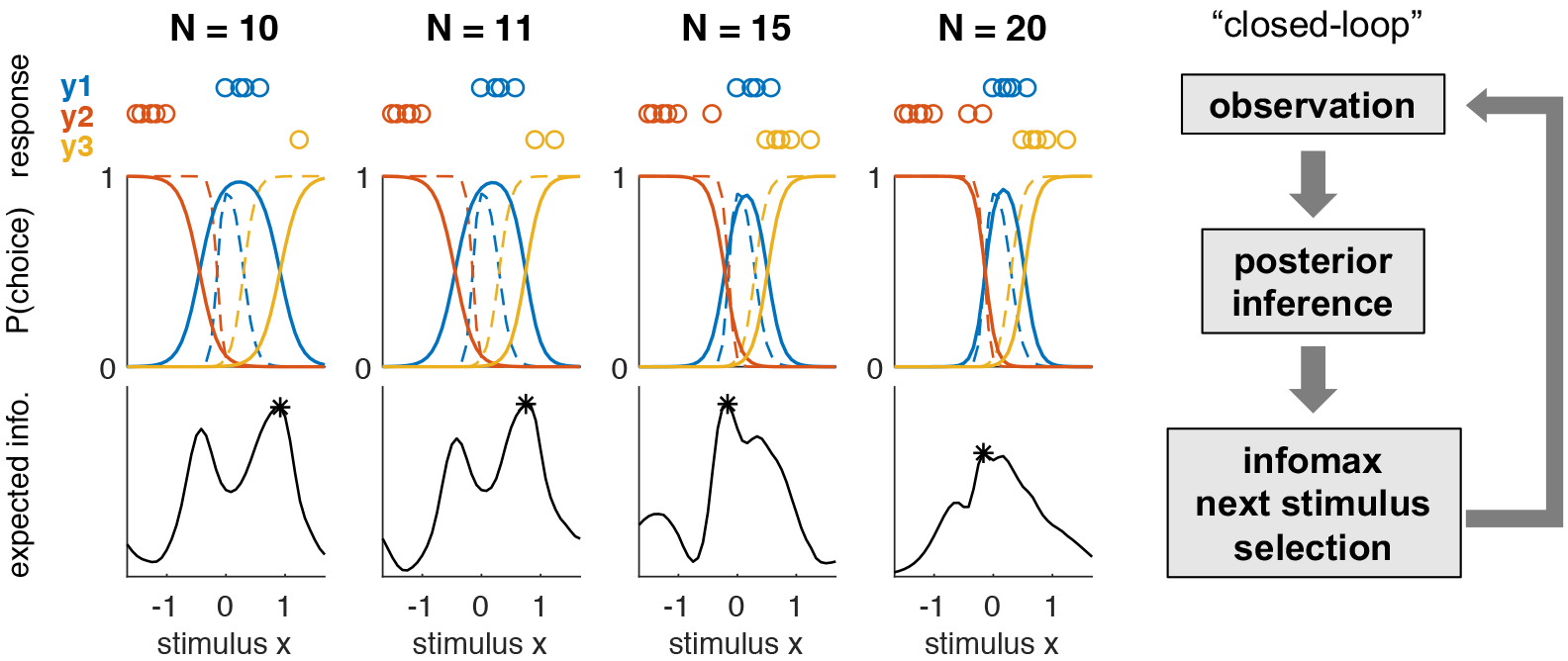
**Example of infomax adaptive stimulus selection,** simulated with a three-alternatives lapse-free model on **1D** stimulus. The figure shows how given a small set of data (the stimulus-response pairs shown in top row), the PFs are estimated based on the accumulated data (middle row), and the next stimulus is chosen to maximize the expected information gain (bottom row). Each column shows the instance after the *N* observations in a single adaptive stimulus selection sequence, for *N* = 10,11,15 and 20 respectively. In the middle row, the estimated PFs (solid lines) quickly approach the true PFs (dashed lines) through the adaptive and optimal selection of stimuli. This example was generated using the Laplace approximation based algorithm, with an independent Gaussian prior over the weights with mean zero and standard deviation *σ* = 10.

Selecting the optimal stimulus thus requires maximizing the mutual information over the set of all possible stimuli {x}. Since each evaluation of the mutual information involves a high-dimensional integral over parameter space and response space, this is a highly computationally demanding task. In the next sections, we present two algorithms for efficient infomax stimulus selection based on each of the two approximate inference methods described previously.

### Infomax with Laplace approximation

Calculation of the mutual information is greatly simplified by a Gaussian approximation of the posterior. The entropy of a Gaussian distribution with covariance *C* is equal to 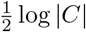 up to a constant factor. If we expand the mutual information as in (eq. 12), and recall that we need only minimize the expected posterior entropy after observing the response, the optimal stimulus for time-step *t* + 1 is given by:

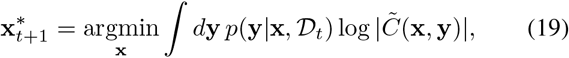

where *C̃*(x, y) is the covariance of the updated (Gaussian) posterior after observing stimulus-response pair (x, y). To evaluate the updated covariance *C̃*(x, y) under the Laplace approximation, we would need to numerically optimize the posterior for ***θ*** for each possible resonse y, for any candidate stimulus x, which would be computationally infeasible. We therefore use a fast approximate method for obtaining a closed-fonn update for *C̃*(x, y) from the current posterior covariance *C*_*t*_, following an approach developed in Lewi et al. (2009). See Appendix C for details. Note that this approximate sequential update is only used for calculating the expected utility of each candidate stimulus by approximating the posterior distribution at the next trial. For obtaining the MAP estimate of the current model parameter, ***θ***_*t*_, numerical optimization needs to be performed using the full accumulated data 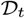 each time.

Once we have log |*C̃*(x, y)| for each given stimulus-observation pair, we numerically sum this over a set of discrete counts y that are likely under the posterior-predictive distribution. This is done in two steps, by separating the integral in (eq. 19) as:

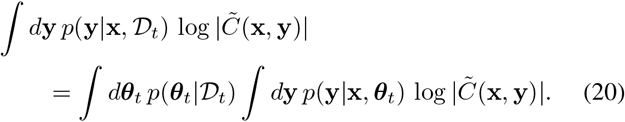

Note that the outer integral is over the current posterior 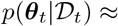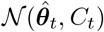 which is to be distinguished from the future posterior 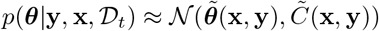 whose entropy we are trying to minimize. Whereas the inner integral is simply a weighted sum over the set of outcomes y, the outer integral over the parameter ***θ*** is in general challenging, especially when the parameter space is high-dimensional. In the case of the standard multinomial logistic model that does not include lapse, we can exploit the linear structure of model to reduce this to a lower-dimensional integral over the space of the linear predictor, which we evaluate numerically using Gauss-Hermite quadrature (Heiss & Winschel, 2008). (This integral is 1D for classic logistic regression, and (*k*−1)-dimensional for multinomial logistic regression with *k* classes; see Appendix C for details.) When the model incorporates lapses, the full parameter vector ***θ*** = [**w**^⊤^, **u**^⊤^]^⊤^ includes the lapse parameters in addition to the weights **w**. In this case, our method with Laplace approximation may suffer from reduced accuracy due to the fact that the posterior may be less closely approximated by a Gaussian.

In order to exploit the convenient structure of reduced integral over the weight space, we choose to maximize the *partial* information between the observation and the psychophysical weights, *I*(w; y|x), instead of the full information *I*(***θ***; y|x). This is a reasonable approximation in many cases where the stimulus-dependent behavior is the primary focus of the psychometric experiment (also see Prins (2013) for a similar approach). However, we note that this is the only piece in this work where we treat the weights separately from the lapse parameters; posterior inference is still performed for the full parameter ***θ***. Thus for Laplace-based infomax exclusively, the partial covariance 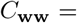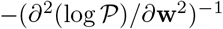 is used in place of the full covariance 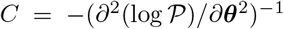, where 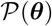 is the posterior distribution over the full parameter space. Because the positive semidefiniteness of the partial covariance is still not guaranteed, it needs to be approximated to the nearest symmetric positive semi-definite matrix when necessary (Higham, 1988). We can show, however, that the partial covariance is asymptotically positive semi-definite in the small-lapse limit (Appendix A).

### Infomax with MCMC

Sampling-based inference provides an attractive alternative to Laplace’s method when the model includes non-zero lapse rates, where the posterior may be less well approximated by a Gaussian. To compute mutual information from samples, it is more convenient to use the expansion given in (eq. 15), so that it is expressed as the expected uncertainty reduction in entropy of the response y, instead of a reduction in the posterior entropy. This will make it straightforward to approximate integrals needed for mutual information by Monte Carlo integrals involving sums over samples. Also note that we are back in the full parameter space; we no longer treat the lapse parameters separately, as we did for the Laplace-based infomax.

Given a set of posterior samples {***θ***_*m*_} from 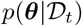, the posterior distribution at time *t*, we can evaluate the mutual information using sums over “potential” terms that we denote by

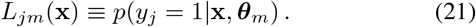

This allows us to evaluate the conditional response entropy as

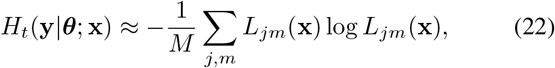

and the marginal response entropy as

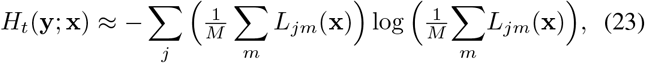

where we have evaluated the posterior-predictive distribution as

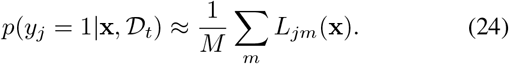

Putting together these terms, the mutual information can be evaluated as

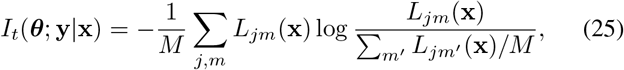

which is straightforward to evaluate for a set of candidate stimuli {x}. The computational cost of this approach is therefore linear in the number of samples, and the primary concern is the cost of obtaining a representative sample from the posterior.

## Results

We consider two approaches for testing the performance of our proposed stimulus-selection algorithms, one using simulated data, and a second using an offline analysis of data from real psychophysical experiments.

### Simulated experiments

We first tested the performance of our algorithms using simulated data from a fixed psychophysical observer model. In these simulations, a stimulus x was selected on each trial and the observer’s response y was sampled from a “true” psychometric function, *p*_true_(**y**|**x**) = *p*(**y**|**x**, ***θ***_true_).

We considered psychophysical models defined on a continuous 2-dimensional stimulus space with 4 discrete response alternatives for every trial, corresponding to the problem of estimating the direction of 2D stimulus moving along one of the four cardinal directions (up, down, left, right). We computed expected information gain over a set of discrete stimulus values corresponding to 21 × 21 square grid (Fig.5A). The stimulus plane is colored in Fig. 5A, to indicate the most likely response (one of the four alternatives) in each stimulus region. Lapse probabilities λ*c*_*i*_ were set to either zero (the “lapse-free” case), or a constant value of 0.05, resulting in a total lapse probability of λ = 0.2 across the four choices (Fig. 5B). We compared performance of our adaptive algorithms with a method that selected a stimulus uniformly at random from the grid on each trial. We observed that the adaptive methods tended to sample more stimuli near the boundaries between colored regions on the stimulus space (Fig. 5C), which led to more efficient estimates of the PF compared to the uniform stimulus selection approach (Fig. 5D). We also confirmed that the posterior entropy of the inferred parameters decrease more rapidly with our adaptive stimulus sampling algorithms, in all cases (Fig. 5E-F). This was expected because our algorithms explicitly attempt to minimize the posterior entropy, by maximizing the mutual information.

For each true model, we compared the performances of four different adaptive methods (Fig. 6A-B), defined by performing inference with MAP or MCMC, and assuming lapse rate to be fixed at zero or including a non-zero lapse parameters. Each of these inference methods was also applied to data selected according to a uniform stimulus selection algorithm. We quantified performance using the mean-squared error (MSE) between the true response probabilities *p*_*ij*_ = *p*(*y* = *j*|x_*i*_, ***θ***_true_) and the estimated probabilities *p̂*_*ij*_ over the 21 × 21 grid of stimulus locations {x_*i*_} and the 4 possible responses {*j*}. For MAP-based inference, estimated probabilities were given by 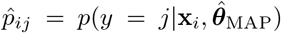. For the MCMC-based inference, probabilities were given by the predictive distribution, evaluated using an average over samples: 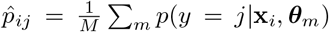, where {***θ***_*m*_} represent samples from the posterior.

When the true model was lapse-free (Fig. 6A), lapse-free and lapse-aware inference methods performed similarly, indicating that there was minimal cost to incorporating parameters governing lapse when lapses were absent. Under all inference methods, infomax stimulus selection outperformed uniform stimulus selection by a substantial margin. For example, infomax algorithms achieved in 50 - 60 trials the error levels that their uniform-stimulus-selection counterparts required 100 trials to achieve.

By contrast, when the true model had a non-zero lapse rate (Fig. 6B), adaptive stimulus selection algorithms based on the lapse-free model failed to select optimal stimuli, performing even worse than uniform stimulus selection algorithms. This emphasizes the impact of model mismatch in adaptive methods, and the importance of a realistic psychometric model. When lapse-aware models were used for inference, on the other hand, both Laplace-based and MCMC-based adaptive stimulus selection algorithms achieved a significant speedup compared to uniform stimulus selection, while MCMC-based adaptive algorithm performed better. This shows that the MCMC-based infomax stimulus selection method can provide an efficient and robust platform for adaptive experiments with realistic models. When the true behavior had lapses, the MCMC-based adaptive stimulus selection algorithm with the lapse-aware model automatically included “easy” trials, which provide maximal information about lapse probabilities. These easy trials are typically in the periphery of the stimulus space (strong-stimulus regimes, referred to as “asymptotic performance intensity” in Prins (2012)).

**Figure 5:**
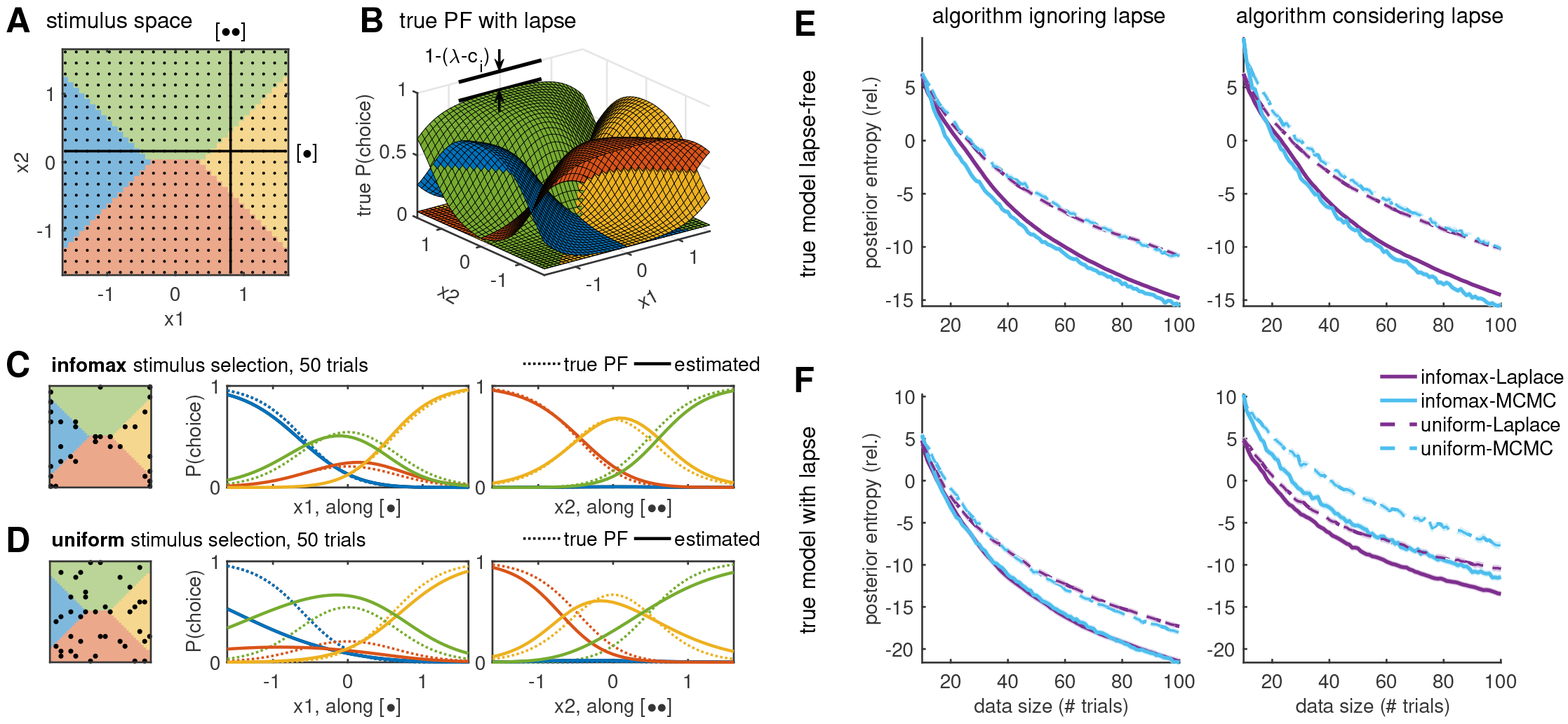
The simulated experiment. **(A)** At each trial, a stimulus was selected from a 2D stimulus plane with a 21 × 21 grid. The two lines, running along *x*_1_ and *x*_2_ respectively, indicate the cross-sections used in **C** and **D** below. Colors indicate the most likely response in the respective stimulus regime, according to the true PF shown in **B**, with a consistent color code. **(B)** Given each stimulus, a simulated response was drawn from a true model with 4 alternatives. Shown here is the model with lapse, characterized by a non-deterministic choice (i.e., the choice probability does not approach 0 or 1) even at an easy stimulus, far from the choice boundaries. **(C-D)** Examples of Laplace-approximation-based inference results after 50 trials, where stimuli was selected either using our adaptive infomax method **(C)** or uniformly **(D)**, as shown on left. In both cases, the true model was lapse-free, and the algorithm assumed that lapse was fixed at zero. The two sets of curves show the cross-sections of the true PF (dotted lines) and the estimated PF (solid lines), along the two lines marked in **A**, after sampling these stimuli. **(E-F)** Traces of posterior entropy from simulated experiments, averaged over 100 runs each. The true model for simulation was either **(E)** lapse-free, or **(F)** with a finite lapse rate of λ = 0.2, with a uniform lapse scenario *c*_*i*_ = 1/4 for each outcome *i* = 1,2, 3,4. In algorithms considering lapse (panels on the right), the shift in posterior entropy is due to the use of partial covariance (with respect to weight) in the case of Laplace approximation. The algorithm either used the classical MNL model that assumes zero lapse (left column), or our extended model that considers lapse (right column). Average performances of adaptive and uniform stimulus selection algorithms are plotted in solid and dashed lines, respectively; Laplace-based and MCMC-based algorithms are plotted in purple and cyan. The lighter lines show standard error intervals over 100 runs, which are very narrow. All sampling-based algorithms used the semi-adaptive MCMC with chain length *M* = 1000.

**Figure 6:**
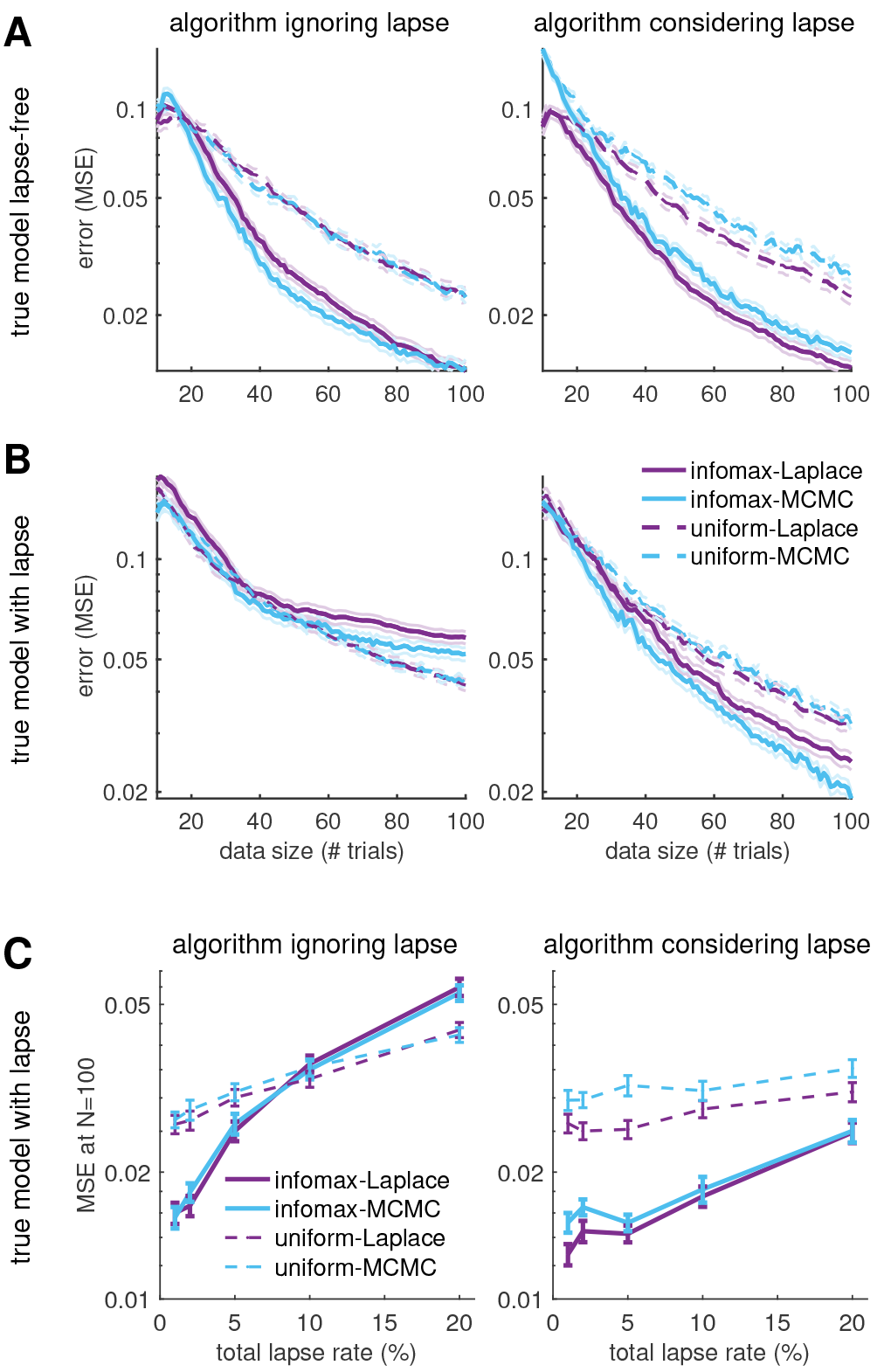
The simulated experiment, continued. We show results from the same set of simulated experiments as in Fig. 5. **(A-B)** Traces of the mean-square error (MSE), where the true model was either **(A)** lapse-free, or **(B)** with a total lapse rate of λ = 0.2, uniformly distributed to each outcome. Standard error intervals are plotted in lighter lines as in Fig.5E-F. **(C)** Effect of lapse, tested by adding varying total lapse rates λ. Shown are the MSE after *N* = 100 trials of each stimulus selection algorithm, equivalent to the endpoints in **B**. Error bars indicate the standard error over 100 runs, equivalent to the lighter-line intervals in the above panels.

However, that the effect of model mismatch due to non-zero lapse only becomes problematic at high enough lapse rate; in the simulation shown in Fig. 5F and Fig. 6B, we used a high lapse rate of λ = 0.2 which is more typical in the case of less sophisticated animals such as rodents (see for example Scott, Constantinople, Erlich, Tank, and Brody (2015)). With lapse rates more typical in well-designed human psychophysics tasks (λ ≲ 0.05; see for example Wichmann and Hill (2001a, 2001b)), infomax algorithms still tend to perform better than uniform sampling algorithms (Fig. 6C).

Finally, we measured the computation time per trial required by our adaptive stimulus selection algorithms on a personal desktop with an Intel i7 processor. With the Laplace-based algorithm, the major computational bottleneck is the parameter space integration in the infomax calculation, which scales directly with the model complexity. We could easily achieve tens-of-milliseconds trials in the case of the simple 2AFC task, and sub-second trials with 2-dimensional stimuli and 4-alternative responses, as used in the current set of simulations (Fig. 7A-B). With the MCMC-based algorithm, the time-per-trial in the sampling-based method is limited by the number of samples in each MCMC chain, *M*, rather than by the model complexity. Using the standard implementation for the Metropolis-Hastings sampler in Matlab, a time-per-trial of ~ 0.1 seconds was achieved with chains shorter than *M* ≲ 200 (Fig. 7C-D, top panels). This length of *M* ≈ 200 was good enough to represent the posterior distributions for our simulated examples (Fig. 7C-D, bottom panels), although we note that longer chains are required to sample a more complex posterior distribution, and this particular length *M* should not be taken as the benchmark in general.

### Optimal re-ordering of real dataset

A second approach for testing the performance of our methods is to perform an off-line analysis of data from real psychophysical experiments. Here we take an existing dataset and use our methods to re-order the trials so that the most-informative stimuli are selected first (also see Lewi, Schneider, Woolley, and Paninski (2011) for a similar approach). To obtain a re-ordering, we iteratively apply our algorithm to the stimuli shown during the experiment. On each trial, we use our adaptive algorithm to select the optimal stimulus from the set of stimuli {x_*i*_} not yet incorporated into the model. This selection takes place without access to the actual responses {y_*i*_}. We then update the posterior using the stimulus x_*i*_ and the response y_*i*_ it actually elicited during the experiment, then proceed to the next trial. We can then ask whether adding the data according to the proposed re-ordering would have led to faster narrowing of the posterior distribution than other orderings.

**Figure 7:**
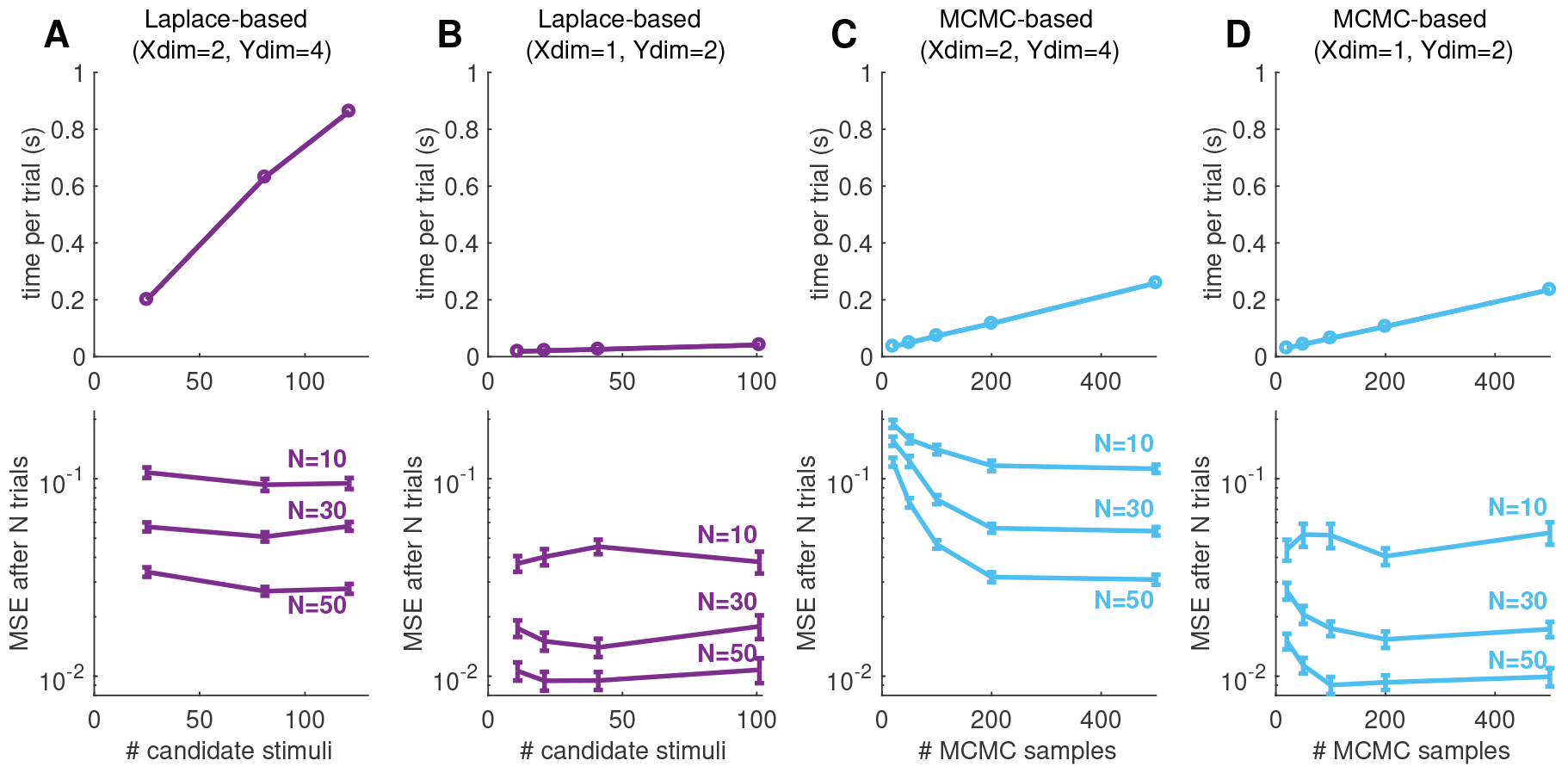
Computation time and accuracy. **(A-B)** The computation times for the Laplace-based algorithms grow linearly with the number of candidate stimulus points, as shown on the top panels, because one needs to perform a numerical integration to compute the expected utility of each stimulus. In general, there is a tradeoff between cost (computation time) and accuracy (inversely related to the estimation error). The bottom panels show the mean-square error of the estimated PF, calculated after completing a sequence of *N* trials, where the 10 initial trials were selected at regular intervals, and the following trials were selected under our adaptive algorithm. Error estimates were averaged over 100 independent sequences. Error bars indicate the standard errors. The true model used were the same as either **(A)** in Fig. 5, with 2-dimensional stimulus and 4-alternative response, described by 9 parameters; or **(B)** in Fig. 3, with 1-dimensional stimulus and binary response, with only 2 parameters (slope and threshold). Different rate at which the computation time increases under the two model reflects the different complexity of numerical quadrature involved. We used lapse-free algorithms in all cases in this example. **(C-D)** We similarly tested the MCMC-based algorithms using the two models as in panels **A-B**. In this case, the computation times (top panels) grow linearly the number of samples in each MCMC chain, and are not sensitive to the dimensionality of the parameter space. On the other hand, the estimation error plots (bottom panels) suggest that a high-dimensional model requires more samples for accurate inference.

To perform this analysis, we used a dataset from macaque monkeys performing a four-alternative motion discrimination task (Churchland, Kiani, & Shadlen, 2008). Monkeys were trained to observe a motion stimulus with dots moving in one of the four cardinal directions, and report this direction of motion with an eye movement. The difficulty of the task was controlled by varying the fraction of coherently moving dots on each trial, with the remaining dots appearing randomly (Fig. 8A). Each moving-dot stimulus in this experiment could be represented as a two-dimensional vector, where the direction of the vector is the direction of the mean movement of the dots, and the amplitude of the vector is given by the fraction of coherently moving dots (a number between 0 and 1). Each stimulus presented in the the experiment was aligned with either one of the two cardinal axes of the stimulus plane (Fig. 8B).

**Figure 8:**
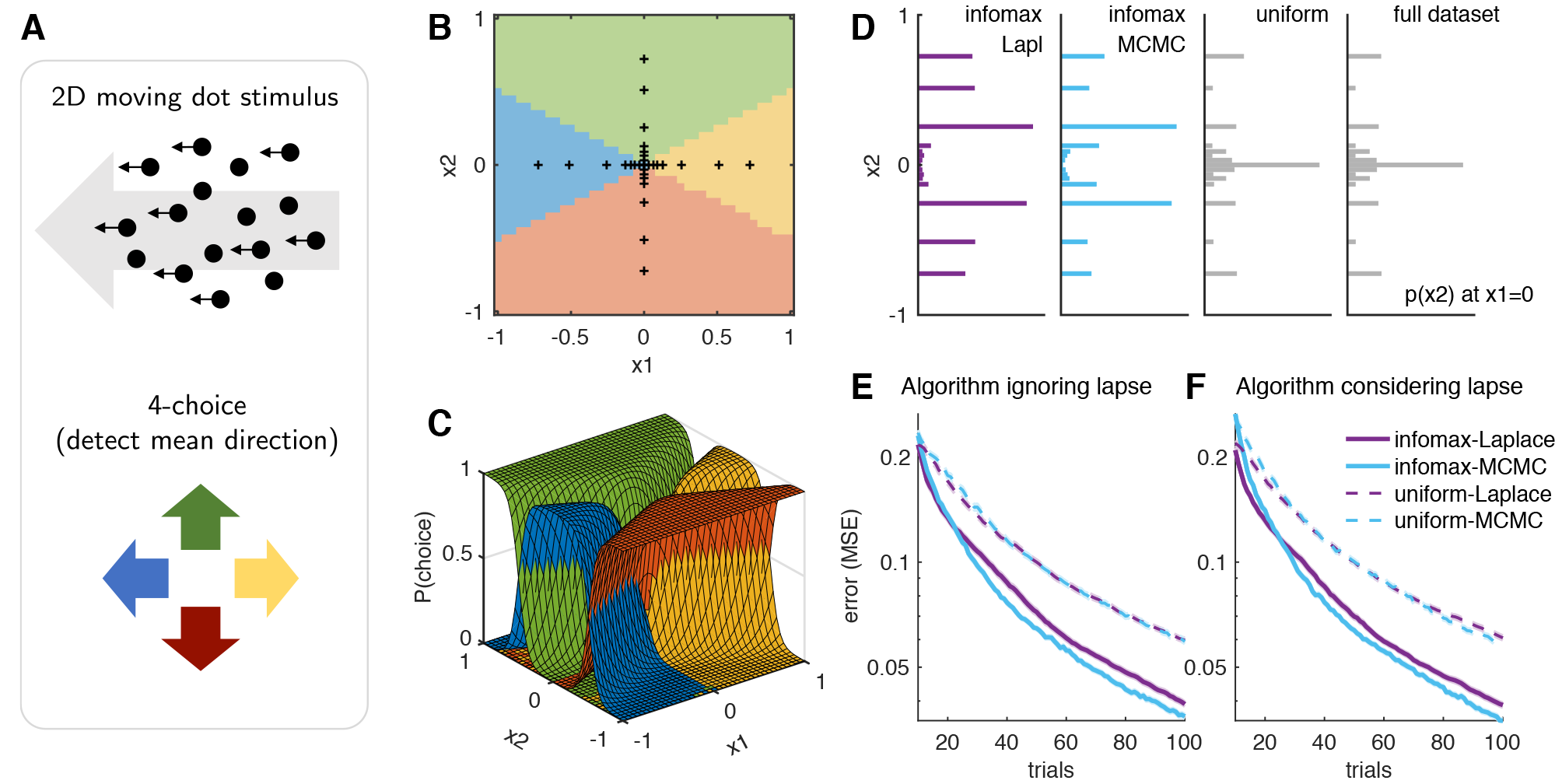
Optimal re-ordering of a real monkey dataset. **(A)** The psychometric task consisted of a 2D stimulus presented as moving dots, characterized by a coherence and a mean direction of movement, and a 4-alternative response. The four choices are color coded consistently in **A**-**C** in this figure. **(B)** The axes-only stimulus space of the original dataset, with 15 fixed stimuli along each axis. Colors indicate the most likely response in the respective stimulus regime according to the best estimate of the PF. **(C)** The best estimate of the PF of monkeys in this task, inferred from all observations in the dataset. **(D)** Stimuli selection in the first *N* = 100 trials during the re-ordering experiment, under the inference method that ignores lapse. Shown are histograms of *x*_2_ along one of the axes, *x*_1_ = 0, averaged over 100 independent runs in each case. **(E-F)** Error traces under different algorithms, averaged over 100 runs. Both Laplace-based (purple) and MCMC-based (cyan; with M = 1000) algorithms achieve significant speedups over uniform sampling. Because the monkeys were almost lapse-free in this task, inference methods that ignore lapse **(E)** and consider lapse **(F)** performed similarly. Standard error intervals over 100 runs are shown in lighter lines, but are very narrow.

The PF for this dataset consists of a set of four 2D curves, where each curve specifies the probability of choosing a particular direction as a function of location in the 2D stimulus plane (Fig. 8C).

This monkey dataset contained more than 10,000 total observations at 29 distinct stimulus conditions, accumulating more than 300 observations per stimulus. This multiplicity of observations per stimulus ensured that the posterior distribution given the full dataset was narrow enough that it could be considered to provide a “ground truth” psychometric function against which the inferences based on the re-ordering experiment could be compared.

The first 100 stimuli selected by the infomax algorithms had noticeably different statistics than the full dataset or its uniform sub-sampling (the first *N* = 100 trials under uniform sampling).

On the other hand, the sets of stimuli selected by both MAP-based and MCMC-based infomax algorithms were similar. Fig. 8D shows the histogram of stimulus component along one of the axes, *p*(*x*_2_ |*x*_1_ = 0), from the first *N* = 100 trials, averaged over 100 independent runs under each stimulus selection algorithm using the lapse-free model.

Because the true PF was unknown, we compared the performance of each algorithm to an estimate of the PF from the entire dataset. When using the MAP algorithm, the full-dataset PF was given by 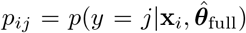, evaluated at the MAP estimate of the log posterior, 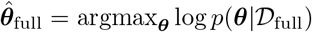, given the full dataset 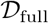. For the MCMC algorithm, the full-dataset PF was computed by 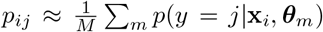, where the MCMC chain 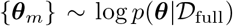 sampled the log posterior given the full dataset. The re-ordering test on the monkey dataset showed that our adaptive stimulus sampling algorithms were able to infer the PF to a given accuracy in a smaller number of observations, compared to a uniform sampling algorithm (Fig. 8E-F). In other words, data collection could have been faster with an optimal reordering of the experimental procedure.

### Exploiting the full stimulus space

In the experimental dataset considered in the previous section, the motion stimuli were restricted to points along the cardinal axes of the 2D stimulus plane (Fig. 8B) (Churchland et al., 2008). In some experimental settings, however, the psychometric functions of interest may lack identifiable axes of alignment or may exhibit asymmetries in shape or orientation. Here we show that in such cases, adaptive stimulus selection methods can benefit from the ability to select points from the full space of possible stimuli.

We performed experiments with a simulated observer governed by the lapse-free psychometric function estimated from the macaque monkey dataset (Fig. 8C). This psychometric function was either aligned to the original stimulus axes (Fig. 9A-B) or rotated counter-clockwise by 45 degrees (Fig. 9C). We tested the performance of adaptive stimulus selection using the Laplace infomax algorithm, with stimuli restricted to points along the cardinal axes (Fig. 9A), or allowed to a grid of points in the full 2D stimulus plane (Fig. 9B-C).

The simulated experiment indeed closely resembled the results of our dataset re-ordering test in terms of the statistics of adaptively selected stimuli (compare Fig. 9A to the purple histogram in Fig. 8D). With the full 2D stimulus space aligned to the cardinal axes, on the other hand, our adaptive infomax algorithm detected and sampled more stimuli near the boundaries between colored regions in the stimulus plane, which were usually not on the cardinal axes (Fig. 9B). Finally, we also observed that this automatic exploitation of the stimulus space was not limited by the lack of alignment between the PF and the stimulus axes; our adaptive infomax algorithm was just as effective in detecting and sampling the boundaries between stimulus regions in the case of the unaligned PF (Fig. 9C).

The error traces in Fig. 9D show that we can infer the PF at a given accuracy in an even fewer number of observations using our adaptive algorithm on the full 2D stimulus plane (orange curves), compared to the cardinal-axes design (black curves). It also confirms that we can infer the PF accurately and effectively with an unaligned stimulus space (red curves), as well as with an aligned stimulus space. For comparison purposes, all errors were calculated over the same 2D stimulus grid, even when the stimulus selection was from the cardinal axes. (This had negligible effects on the resulting error values: compare the black curves in Fig. 9D and the purple curves in Fig. 8E.)

## Discussion

We developed effective Bayesian adaptive stimulus selection algorithms for inferring psychometric functions, with an objective of maximizing the expected informativeness of each stimulus. The algorithms select an optimal stimulus adaptively in each trial, based on the posterior distribution of model parameters inferred from the accumulating set of past observations.

We emphasized that in psychometric experiments, especially with animals, it is crucial to use models that can account for the non-ideal yet common behaviors, such as omission (no response; an additional possibility for the outcome) or lapse (resulting in a random, stimulus-independent response). Specifically, we constructed a hierarchical extension of a multinomial logistic (MNL) model that incorporates both omission and lapse. Although we did not apply these additional features to real data, we performed simulated experiments to investigate their impacts on the accurate inference of psychometric functions. To ensure applicability of the extended model in real-time closed-loop adaptive stimulus selection algorithms, we also developed efficient methods for inferring the posterior distribution of the model parameters, with approximations specifically suited for sequential experiments.

**Figure 9:**
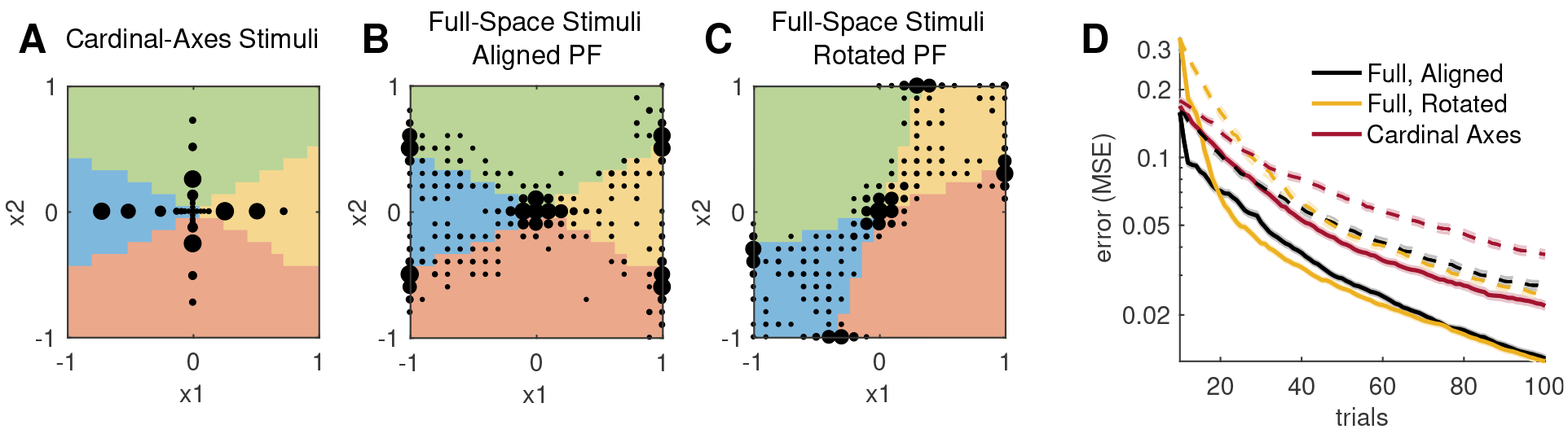
Design of multi-dimensional stimulus space. **(A-C)** Three different stimulus space designs were used in a simulated psychometric experiment. Responses were simulated according to fixed lapse-free PFs, matched to our best estimate of the monkey PF (Fig. 8C). Stimuli were selected within the respective stimulus spaces, **(A)** the cardinal-axes design, as in the original experiment; **(B)** full stimulus plane, with the PF aligned to the cardinal axes of the original stimulus space; **(C)** full stimulus plane, with rotated PF. The black dots in **A-C** indicate which stimuli were sampled by the Laplace-based infomax algorithm during the first *N* = 100 trials of simulation, where the dot size is proportional to the number of trials in which each stimulus was selected (averaged over 20 independent runs, and excluding the 10 fixed initial stimuli). **(D)** The corresponding error traces, under infomax (solid lines) or uniform (dashed lines) stimulus selection, averaged over 100 runs respectively. Colors indicate the three stimulus space designs, as shown in **A**-**C**. Standard error intervals over 100 runs are shown in lighter lines.

### Advantages of adaptive stimulus selection

We observed two important advantages of using Bayesian adaptive stimulus selection methods in psychometric experiments. First, we showed that our adaptive stimulus selection algorithms achieved significant speed-ups in learning time (number of measurements), both on simulated data and in re-ordering test of a real experimental dataset, with and without lapse in the underlying behavior. Importantly, the success of the algorithm depends heavily on the use of the correct model family; for example, adaptive stimulus selection fails when a classical (lapse-ignorant) model was used to measure behavior with a finite lapse rate. Based on the simulation results, it seems good practice to always use the lapse-aware model unless the behavior under study is known to be completely lapse-free, although it should be checked that the addition of the lapse parameters does not make the inference problem intractable, given the constraints of the specific experiments. (One way to check this is using a simulated experiment, where lapse is added to the psychometric function inferred by lapse-free model; similarly to what we did in this paper.) The computational cost for incorporating lapses amounts to having *k* additional parameters to sample, one per each available choice, which is independent from the dimensionality of the stimulus space.

Second, we demonstrated that our adaptive stimulus selection study has implications on the optimization of the experimental designs more generally. Contrary to the conventional practice of accumulating repeated observations at a small set of fixed stimuli, we suggest that the (potentially high-dimensional) stimulus space can be exploited more efficiently using our Bayesian adaptive stimulus selection algorithm. Specifically, the adaptive stimulus selection algorithm can automatically detect the structure of the stimulus space (with respect to the psychometric function) as part of the process. We also showed that there are benefits of using the full stimulus space even when the PF is aligned to the cardinal axes of the stimulus space.

### Comparison of the two algorithms

Our adaptive stimulus selection algorithms were developed based on two methods for effective posterior inference: one based on local Gaussian approximation (Laplace approximation) of the posterior, and another based on MCMC sampling. The well-studied analytical method based on the Laplace approximation is fast and effective in simple cases, but becomes heavier in the case of more complicated PFs, because the computational bottleneck is the numerical integration over the parameter space that needs to be performed separately for each candidate stimulus. In the case of sampling-based methods, on the other hand, the computational speed is constrained by the number of MCMC samples used to approximate the posterior distribution, but not directly by the number of parameters or the number of candidate stimuli. In general, however, accurately inferring a higher-dimensional posterior distribution requires more samples, and therefore a longer computation time. We note that our semi-adaptive turning algorithm helps with the cost-accuracy tradeoff by optimizing the sampling accuracy in a given number of samples, without human intervention. although it does not reduce the computation time itself.

To summarize, when the PF under study is low-dimensional and well-described by the multinomial logistic model, for example in a 2AFC study with human subjects, Laplace-based approach provides a lightweight and elegant approach. But if the PF is higher-dimensional or deviates significantly from the ideal model (e.g., large lapse), MCMC sampling provides a flexible and affordable solution. Results suggest that our MCMC-based algorithm will be applicable to most animal psychometric experiments, as the model complexities are not expected to significantly exceed our simulated example. However, one should always make sure that the number of MCMC samples being used is sufficient to sample the posterior distribution under study.

### Limitations and Open Problems

One potential drawback of adaptive experiments is the undesired possibility that the psychometric function of the observer might adapt to the distribution of stimuli presented during the experiments. If this is the case, the system under measurement would no longer be stationary, nor independent of the experimental design, profoundly altering the problem one should try to solve. The usual assumption in psychometric experiments is that well trained observers exhibit stationary behavior on the timescale of an experiment; under this assumption, the order of data collection cannot bias inference MacKay (1992). However, the empirical validity of this claim remains a topic for future research.

One approach for mitigating non-stationarity is to add regressors to account for the history dependence of psychophysical behavior. Recent work has shown that extending a psychophysical model to incorporate past rewards (Bak et al., 2016; Busse et al., 2011; Cor-rado, Sugrue, Seung, & Newsome, 2005; Lau & Glimcher, 2005), past stimuli (Akrami, Kopec, Diamond, & Brody, 2018) or the full stimulus-response history (Friind, Wichmann, & Macke, 2014) can provide a more accurate description of the factors influencing responses on a trial-by-trial basis.

Our work leaves open a variety of directions for future research. One simple idea is to re-analyze old datasets under the multinomial response model with omissions included as a separate response category; this will reveal whether omissions exhibit stimulus dependence (e.g., occurring more often on difficult trials), and provide greater insight into the factors influencing psychophysical behavior on single trials. Another set of directions is to extend the multinomial logistic observer model to obtain a more accurate or more flexible model of psychophysical behavior; particular directions include models with nonlinear stimulus dependencies or interaction terms (Cowley, Williamson, Clemens, Smith, & Byron, 2017; DiMattina & Zhang, 2011; Hyafil & Moreno-Bote, 2017; Neri & Heeger, 2002), models with output nonlinearities other than the logistic (Kontsevich & Tyler, 1999; Schütt et al., 2016; A. B. Watson, 2017; A. B. Watson & Pelli, 1983), or models that capture overdispersion, e.g., due to non-stationarities of the observer, via a hierarchical prior (Schütt et al., 2016). In general, such extensions will be much easier to implement with the MCMC-based inference method, due to the fact that it does not rely on gradients or Hessians of a particular parametrization of log-likelihood. Finally, it may be useful to consider the same observer model under optimality criteria other than mutual information — recent work has shown that infomax methods do not necessarily attain optimal performance according to alternate metrics (e.g., mean squared error, I. M. Park and Pillow (2017); M. Park et al. (2014)) — or using non-greedy selection criteria that optimize stimulus selection based on a time horizon longer than the next trial (Kim et al., 2017; King-Smith et al., 1994).

## Code availability

A Matlab implementation of our methods is available online at https://github.com/pillowlab/adaptivePsychophysicsToolbox.

## Acknowledgements

We thank Anne Churchland for sharing the monkey data. JHB was supported by the Samsung Scholarship for the study at Princeton. JWP was supported by grants from the McKnight Foundation, Simons Collaboration on the Global Brain (SCGB AWD1004351) and the NSF CAREER Award (IIS-1150186). Computational work was performed using resources at Princeton University and the KIAS Center for Advanced Computing.

## Appendix A

**Log likelihood for the classical MNL.** Here we provide more details about the log likelihood *L* = **y**^⊤^ log **p** under the multinomial logistic model (6), first in the lapse-free case.

A convenient property of the multinomial logistic model (a property common to all generalized linear models) is that the parameter vector *p*_*i*_ governing *y* depends only on a 1-dimensional projection of the input, *V*_*i*_ = *ϕ*^⊤^**w**_*i*_, which is known as the *linear predictor*. Recall that *ϕ* = *ϕ*(x) is the input feature vector. In the multinomial case, it is useful to consider the column vector of linear predictors for a single trial, **V** = [*V*_1_, · · ·, *V*_*k*_]^⊤^, and the concatenated weight vector 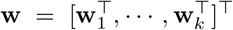, consisting of all weights stacked into a single vector. We can summarize their linear relationship as **V** = *X***w**, where *X* is a block diagonal matrix containing *k* blocks of *ϕ*^⊤^ along the diagonal. In other words,

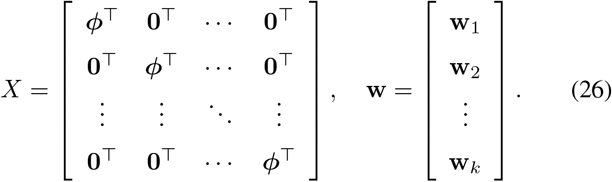

***Derivatives.*** It is convenient to work in terms of the linear predictor **V** = {*V*_*i*_} first. If *N*_*y*_ ≡ ∑_*i*_ *y*_*i*_ = 1 is the total number of responses per trial, the first and second derivatives of *L* with respect to **V** are ∂*L*/∂*V*_*j*_ = *y*_*j*_−*N*_*y*_*p*_*j*_ and ∂^2^*L*/∂*V*_*i*_∂*V*_*j*_ = *N*_*y*_*p*_*i*_(*δ*_*ij*_−*p*_*j*_), respectively. Rewriting in vector forms, we have

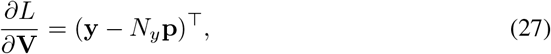

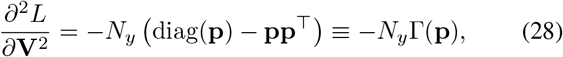

where diag(**p**) = [*p*_*i*_*δ*_*ij*_] is a square matrix with the elements of **p** on the diagonal, and zeros otherwise.

Putting back in terms of the weight vector **w** is easy, thanks to the linear relationship **V** = *X***w**:

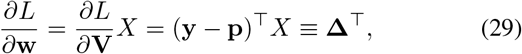

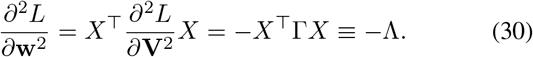

***Concavity.*** Importantly, *L* is concave with respect to **V** (and therefore with respect to **w**). To prove the concavity of *L*, we show that the Hessian *H* = −diag(**p**) + **pp**^⊤^ = −Γ is negative semi-definite, which is equivalent to showing that **z**^⊤^Γ**z** ≥ 0:

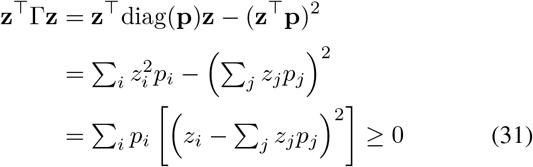

for an arbitrary vector **z**.

**Log likelihood with lapse.** With a finite lapse rate λ, to recap, the multinomial logistic model is modified as *p*_*i*_ = (1 - λ)*q*_*i*_ + λ*c*_*i*_ where

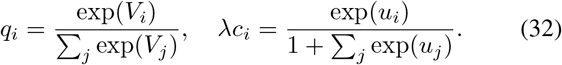

Let us introduce the following abbreviations,

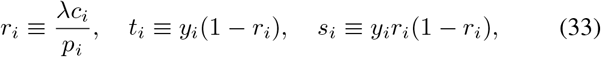

where the dimensionless ratio *r* ∊ [0,1] can be considered as the order parameter for the effect of lapse.

***Derivatives with respect to the weights.*** Differentiating with the linear predictor **V**, we get

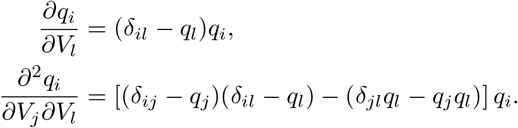

which leads to

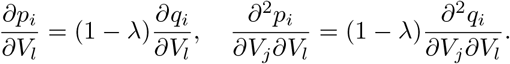

We are interested in the derivatives of the log likelihood *L* = **y**^⊤^ log **p** with respect to **V**. The partial gradient:

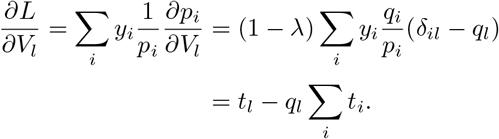

Similarly, the partial Hessian is written as

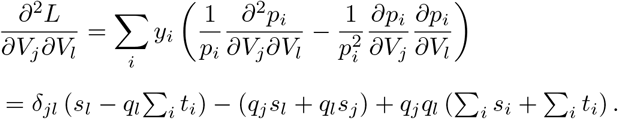

In vector forms, and with τ ≡ ∑_*i*_*t*_*i*_ and *σ* ≡ ∑_*i*_*s*_*i*_

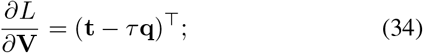

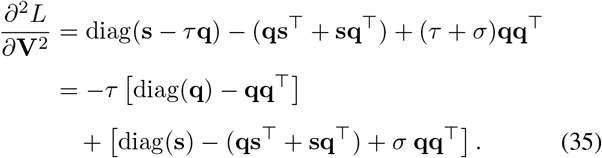

Note that we recover *t*_*i*_ → *y*_*i*_ and *s*_*i*_ → 0 in the lapse-free limit λ → 0. Hence the first square bracket in (35) reduces back to the lapse-free Hessian, while the second square bracket vanishes as λ → 0.

In the presence of lapse, one might still be interested in the partial Hessian with respect to the weight parameters, *H* ≡ ∂^2^*L*/∂**V**^2^, which should be evaluated as in (35). To test the negative semi-definiteness of this partial Hessian, again for an arbitrary vector **z**, we end up with

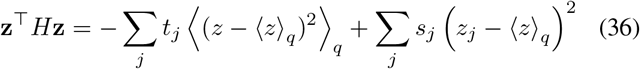

where 〈*x*〉_*q*_ = ∑_*j*_*x*_*j*_*q*_*j*_. The partial Hessian is asymptotically negative semi-definite (which is equivalent to the log likelihood being concave) in the lapse-free limit, where *t*_*j*_ → *y*_*j*_ and *s*_*j*_ → 0.

***Derivatives with respect to lapse parameters.*** From (2) and (3), we have *p*_*i*_ = (1 − λ)*q*_*i*_ + λ*c*_*i*_ where

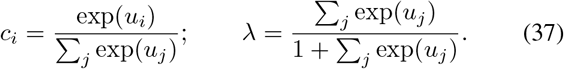

Differentiating with respect to the auxiliary lapse parameter *u*_*i*_,

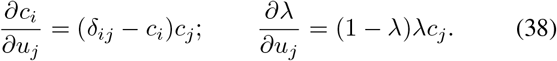

The gradient is then

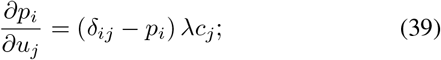

using the abbreviations in (33), the gradient of the log likelihood is

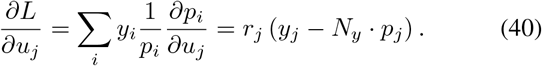

Second derivative with respect to lapse:

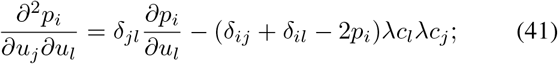

it is useful to notice that

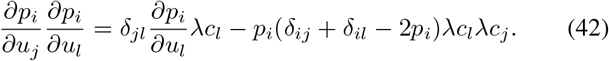

The corresponding part of the Hessian:

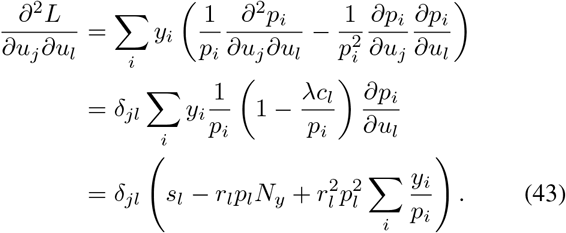

Finally, the mixed derivative:

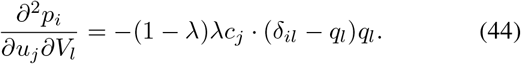

again it is useful to notice that

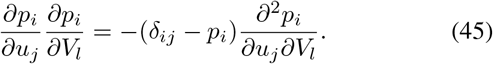

Hence

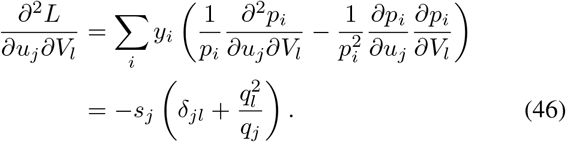

From (40), (43) and (46), we see that all derivatives involving the lapse parameter scale with at least one order of *r*, therefore vanishing in the lapse-free limit λ → 0.

## Appendix B

**The Metropolis-Hastings algorithm.** The Metropolis-Hastings algorithm (Metropolis et al., 1953) generates a chain of samples, using a proposal density and a method to accept or reject the proposed moves.

A proposal is made at each iteration, where the algorithm randomly chooses a candidate for the next sample value x′ based on the current sample value x_*t*_. The choice follows the proposal density function, x′ ~ *Q*(x′|x_*t*_). When the proposal density *Q* is symmetric, for example a Gaussian, the sequence of samples is a random walk. In general the width of *Q* should match with the statistics of the distribution being sampled, and individual dimensions in the sampling space may behave differently in the multivariate case; finding the appropriate *Q* can be difficult.

The proposed move is either accepted or rejected with some probability; if rejected, the current sample value is reused in the next iteration, x′ = x_*t*_. The probability of acceptance is determined by comparing the values of *P*(x_*t*_) and *P*(x′), where *P*(x) is the distribution being sampled. Because the algorithm only considers the acceptance ratio *ρ* = *P*(x′)/*P*(x_*t*_) = *f* (x′)/*f*(x_*t*_) where *f*(x) can be any function proportional to the desired distribution *P*(x), there is no need to worry about the proper normalization of the probability distribution. If *ρ* ≥ 1, the move is always accepted; if *ρ* < 1, it is accepted with a probability *ρ*. Consequently the samples tend to stay in the high-density regions, visiting the low-density regions only occasionally.

**Optimizing the sampler.** One of the major difficulties in using the MCMC method is to make an appropriate choice of the proposal distribution, which may significantly affect the performance of the sampler. If the proposal distribution is too narrow, it will take a long time for the chain to diffuse away from the starting point, producing a chain with highly correlated samples, requiring a long time to achieve independent samples. On the other hand if the proposal distribution is too wide, most of the proposed moves would be rejected, once again resulting in the chain stuck at the initial point. In either case the chain would “mix” poorly (Rosenthal, 2011). In this paper we restrict our consideration to the Metropolis-Hastings algorithm (Metropolis et al., 1953), although the issue of proposal distribution optimization is universal in most variants of MCMC algorithms, only with implementation-level differences.

The basic idea is that the optimal width of the proposal distribu-tion would be determined in proportion to the typical length scale of the distribution being sampled. This idea was made precise in the case of a stationary random-walk Metropolis algorithm with Gaussian proposal distributions, by comparing the covariance matrix Σ_*p*_ of the proposal distribution to the covariance matrix Σ of the sampled chain. Once a linear scaling relation Σ_*p*_ = *s*_*d*_Σ is fixed, it was observed that it is optimal to have *s*_*d*_ = (2.38)^2^/*d* where *d* is the dimensionality of the sampling space (Gelman et al., 1996; Roberts et al., 1997). An adaptive Metropolis algorithm (Haario et al., 2001) followed this observation, where the Gaussian proposal distribution adapts continuously as the sampling progresses. Their adaptive algorithm used the same scaling rule Σ_*p*_ = *s*_*d*_Σ, but updates Σ_*p*_ at each proposal where Σ is covariance of the samples accumulated so far. Additionally, a small diagonal component was added for stability, as Σ_*p*_ = *s*_*d*_(Σ + *ϵI*). We used *ϵ* = 0.0001 in this work.

Here we propose and use the semi-adaptive Metropolis-Hastings algorithm, which is a coarse-grained version of the original adaptive algorithm by Haario et al. (2001). The major difference in our algorithm is that the adjustment of the proposal distribution is made only at the end of each (sequential) chain, rather than at each proposal within the chain. This coarse-graining is a reasonable approximation because we will be sampling the posterior distribution many times as it refines over the course of data collection, once after each trial. Assuming that the change in posterior distribution after each new observation is small enough, we can justify our use of the statistics of the previous chain to adjust the properties of the current chain. Unlike in the fully adaptive algorithm where the proposal distribution needs to stabilize quickly within a single chain, we can allow multiple chains until stabilization, usually a few initial observations - leaving some room for the coarse-grained approximation. This is because, for our purpose, it is not imperative that we have a good sampling of the distribution at the very early stage of the learning sequence where the accuracy is already limited by the smallness of the dataset.

**Figure 10:**
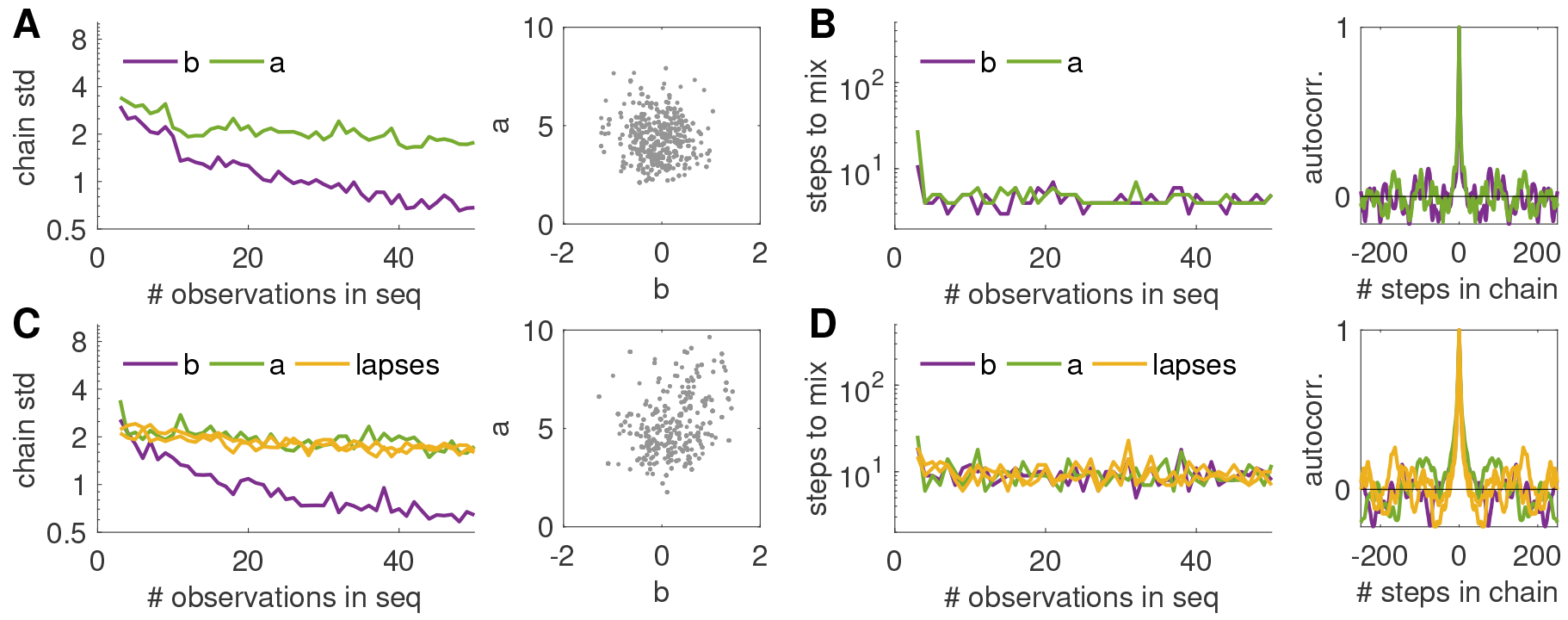
Statistics of the semi-adaptive MCMC in a simulated experiment, with *M* = 1000 samples per chain. We used the same binomial model as in Fig. 3, and the uniform stimulus selection algorithm. **(A-B)** In a lapse-free model: **(A)** The standard deviation of the samples, along each dimension of the parameter space, decreases as the learning progresses, as expected because the posterior distribution should narrow down as more observations are collected. Also shown is the scatter plot of all 1000 samples at the last trial *N* = 50, where the true parameter values are (*a, b*) = (5; 0). **(B)** The mixing time of the chain (number of steps before the autocorrelation falls to 1/*e*) quickly converges to some small value, meaning that the sampler is quickly optimized. Autocorrelation function at the last trial *N* = 50 is shown. **(C-D)** Same information as (A) and (B), but with a lapse rate of λ = 0:1, with uniform lapse (*c*_1_ = *c*_2_ = 1/2).

When applied to the sequential learning algorithm, our semi-adaptive Metropolis sampler shows a consistent well-mixing property after a few initial adjustments, with the standard deviation of each sampling dimension decreasing stably as data accumulate (Fig. 10). Whereas Kujala and Lukka (2006) also had the idea of adjusting the proposal density between trials, their scaling factor was fixed and independent of the sampling dimension. Building on more precise statistical observations, our method generalize well to high-dimensional parameter spaces, typical for multiple-alternative models. Our semi-adaptive sampler provides an efficient and robust alternative to the particle filter implementations (Kujala & Lukka, 2006), which has the known problem of weight degeneration (DiMattina, 2015) as the posterior distribution narrows down with the accumulation of data.

## Appendix C

**Fast sequential update of the posterior, with Laplace approximation.** Use of Laplace approximation was shown to be particularly useful in a sequential experiment (Lewi et al., 2009), where it can be assumed that the posterior distribution after the next trial in sequence, 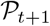, would not be very different from the current posterior 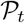. Let us consider the lapse-free case ***θ*** = **w** for the moment, where the use of Laplace approximation is valid. Rearranging from (7) and (9), the sequential update for the posterior distribution is

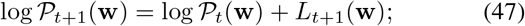

or with Laplace approximation,

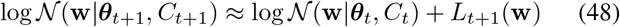

where *L*_*i*_(**w**) = log*p*(**y**_*i*_|**x**_*i*_, **w**) is a shorthand for the log likelihood of the *i*-th observation.

With this, we can achieve a fast sequential update of the posterior without performing the full numerical optimization each time. Because the new posterior mode ***θ***_*t*+1_ is where the gradient vanishes, it can be approximated from the previous mode ***θ***_*t*_ by taking the first derivative of (48). The posterior covariance *C*_*t*+1_ is similarly approximated by taking the second derivate.

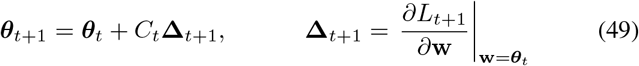

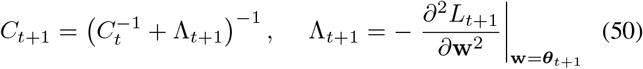

Using the matrix inversion lemma (Henderson & Searle, 1981), we can rewrite the posterior covariance update as

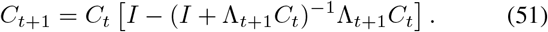

Unlike in the earlier application of this trick (Lewi et al., 2009), the covariance matrix update (50) is not a rank-one update, because of the multinomial nature of our model (our linear predictor y is a vector, not a scalar as in a binary model).

**Integration over the parameter space: reducing the integration space.** The evaluation of expected utility function usually involves a potentially high-dimensional integral over the parameter space. With the Gaussian approximation of the posterior, we can reduce and standardize the integration space. The process consists of three steps: diagonalization, marginalization, and standardization. First we choose a new “coordinate system” of the (say *q*-dimensional) weight space, such that the first *k* elements of the extended weight vector w are coupled one-to-one to the elements of *k*-vector **y**. Then we marginalize to integrate out the remaining (*q* − *k*) dimensions, effectively changing the integration variable from **w** to **y**. Finally, we use Cholesky decomposition to standardize the normal distribution which is the posterior on **y**. The resulting integral is still multi-dimensional, due to the multinomial nature of our model. But once the distribution is standardized, there are a number of efficient numerical integration methods that can be applied. For example, in this work, we use the Sparse Grid method (Heiss & Winschel, 2008) based on Gauss-Hermite quadrature.

***Diagonalization.*** It is clear from (19–20) and (29–30) that all parameter-dependence in our integrand is in terms of the linear predictor **y** = *X***w**. That is, we are dealing with the integral of the form

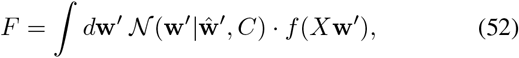

where *C* is the covariance matrix, and 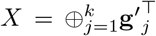 is a fixed matrix constructed from direct sum of *k* vectors. It helps to work in a diagonalized coordinate system, so that we can separate out the relevant dimensions of **w**. We use the singular value decomposition of the design matrix (*X* = *UGV*^⊤^ with *U* = *I* and *V* = *Q*^⊤^). Because of the direct-sum construction, *XX*^⊤^ is already diagonal, and the left singular matrix is always *I* in this case. Then

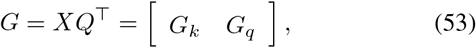

where *G*_*k*_ is a *k* × *k* diagonal matrix and *G*_*q*_ is a *k* × (*q* − *k*) matrix of zeros. We can now denote **w**_*k*_ = (*w*_1_, · · ·, *w*_*k*_) an **w**_*q*_ = (*w*_*k*+1_, · · ·, *w*_*q*_) in the diagonalized variable **w** = *Q***w**′ such that

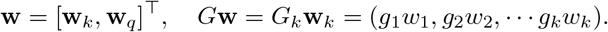

***Marginalization.*** Now we have

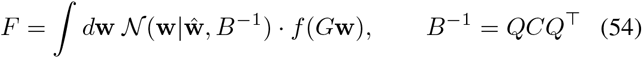

where *B* is the inverse of the *new* covariance matrix after diagonalization. If we block-decompose this matrix,

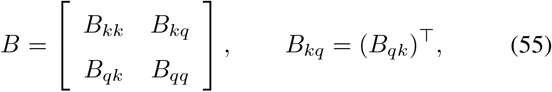

the Gaussian distribution is also decomposed as

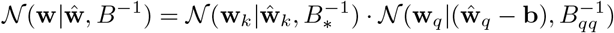

where 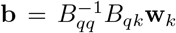 and 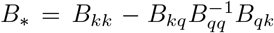. As the non-parallel part **w**_*q*_ is integrated out, we have marginalized the integral. It is useful to recall that if a variable 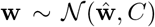 is Gaussian distributed, its linear transform **y** = *X***w** is also Gaussian distributed as 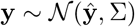, with **ŷ** = *X***ŵ** and Σ = *XCX*^⊤^. Changing the integration variable to **y** = *G*_*k*_ **w**_*k*_ is then straightforward:

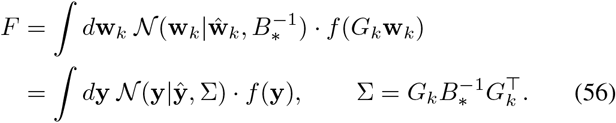

***Standardization.*** Finally, in order to deal with the numerical integration, it is convenient to have the normal distribution standardized. We can use the Cholesky decomposition for the covariance matrix,

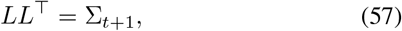

such that the new variable ***θ*** = *L*^−1^(**y** − **ŷ**_*t*+1_) is standard normal distributed. From the above formulation, *L* can be written directly in terms of the Cholesky decomposition of *B*_*_:

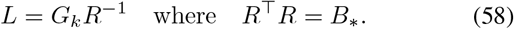

Importantly, with this transformation, each dimension of ***θ*** is independently and identically distributed. The objective function to be evaluated is now

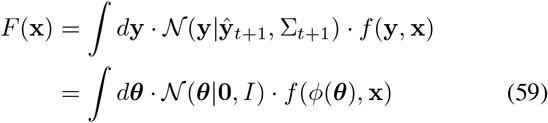

where *ϕ*(***θ***) = **ŷ**_*t*+1_ + *L****θ***. Once the integration is standardized this way, there are a number of efficient numerical methods that can be applied.

